# Low memory T cells blood counts and high naïve regulatory T cells percentage at relapsing remitting multiple sclerosis diagnosis

**DOI:** 10.1101/2022.03.07.483243

**Authors:** João Canto-Gomes, Carolina S. Silva, Rita Rb-Silva, Daniela Boleixa, Ana Martins da Silva, Rémi Cheynier, Patrício Costa, Inés González-Suárez, Margarida Correia-Neves, João Cerqueira, Claudia Nobrega

## Abstract

**Objective:** To assess the peripheral immune system of newly diagnosed relapsing remitting multiple sclerosis (RRMS) patients and compare it to healthy controls (HC).

**Methods:** Cross-sectional study with 30 treatment-naïve newly diagnosed RRMS patients, and 33 sex and age-matched HC. Their peripheral blood mononuclear cells were analysed regarding: i) thymic function surrogates [T cell receptor excision circles (TRECs) and recent thymic emigrants (RTEs)]; ii) naïve and memory CD4^+^ and CD8^+^ T cells subsets; iii) T helper (Th) phenotype and chemokine receptors expression on T cells subsets; iv) regulatory T cell (Tregs) phenotype; and vi) expression of activating/inhibitory receptors by natural killer (NK) and NKT cells. Analyses were controlled for age, sex, and human cytomegalovirus (HCMV) IgG seroprevalence.

**Results:** Newly diagnosed RRMS patients and HC have equivalent thymic function as determined by similar numbers of RTEs, and levels of sjTRECs, DJβTRECs and sj/DJβTREC ratio. In the CD8^+^ T cells compartment RRMS patients have a higher naïve/memory ratio and lower memory cell counts in blood, specifically of effector memory and TemRA CD8^+^ T cells. Among CD4^+^ T cells lower blood counts of effector memory cells are found in patients upon controlling for sex, age and HCMV IgG seroprevalence. RRMS patients have higher percentage of naive Tregs comparing to HC. Percentages of immature CD56^bright^ NK cells expressing the inhibitory receptor KLRG1, and of mature CD56^dim^CD57^+^ NK cells expressing NKp30 are higher in patients. No major alterations are observed on NKT cells. MS severity and time from relapse correlate with immune cells alterations.

**Conclusion:** Characterization of the peripheral immune system of treatment-naïve newly diagnosed RRMS patients unveiled immune features present at clinical onset including lower memory T cells blood counts, particularly among CD8^+^ T cells, higher percentage of naïve Tregs and altered percentages of NK cells subsets expressing inhibitory or activating receptors. These findings might set the basis to better understand disease pathogenesis.

## INTRODUCTION

The immune system has the tricky task of fighting pathogens and tumour cells, while maintaining tolerance to self, avoiding autoimmunity, and ensuring immune homeostasis. T cells are suggested to underlie multiple sclerosis (MS) pathogenesis most probably due to impaired tolerance to myelin-producing cells.^1^ MS is a demyelinating disease clinically characterized by motor and cognitive impairment. The most common MS form (∼85%), the relapsing-remitting MS (RRMS), is characterized by bouts of disability (called relapses) interleaved by periods of recovery.^1^

In a mouse model of MS (experimental autoimmune encephalomyelitis, EAE), peripheral administration of myelin-peptides triggers myelin-specific T cell responses in the central nervous system (CNS) and consequent demyelination.^2^ In humans, the relevance of T cells to MS pathophysiology is highlighted by the efficacy of MS therapies targeting T cells.^3^ Among the T cells involved in MS, the T helper (Th) cells Th1 and Th17 have been proposed as key drivers of the immune attack against cells producing myelin peptides.^4^ These cells are not only characterized by their profile of cytokine production, but also by the differential expression of chemokine receptors that confer them the migratory capacity to home to tissues.^5^ Yet, increasing evidence show the involvement of other T cells in MS, including the CD8^+^ T cells, which outnumber CD4^+^ T cells in MS lesions.^6^ In comparison to healthy controls (HC), MS patients have been suggested to present lower thymic function and thus lower production of new T cells (recent thymic emigrants; RTEs), and an aged peripheral adaptive immune system characterized by an accumulation of memory over naïve T cells.^7–16^

Regulatory T cells (Tregs) are known for inducing tolerance by preventing activation of potential self-reactive T cells.^17^ Tregs have been extensively studied in MS yet observations on percentage, number and function of Tregs in MS are not consensual.^7,12,13,18–20^ Several hypotheses on the ineffectiveness of Tregs in fully preventing MS have been proposed, including their limited capacity to access MS lesions in the CNS (in comparison to conventional T cells), ability of self-reactive T cells to withstand Tregs suppression, and lower number, percentage and/or suppressive capacity of Tregs.^21,22^ Both innate natural killer (NK) and NKT cells are also known to induce tolerance due to their killing potential of self-reactive T cells.^23^ NK cells’ activation and cytotoxicity towards anomalous cells is driven by the lack of engaging of inhibitory receptors and/or up-regulation and engaging of activating receptors to their specific ligands on target cells.^24–26^ NK cells from MS patients were described to have reduced capacity to suppress autologous T cells proliferation *in vitro* in comparison to the ones from HC.^23,26^ NKT cells establish a bridge between innate and adaptive immune system presenting NK and T cell receptors. In MS patients NKT cells were described to be hypo-responsive after stimulation with myelin-derived lipids.^27^

Several immune system alterations have been described in MS patients, however most of those studies lack data at clinical disease onset, before treatment initiation, which hampers the clarification of the immune features that are specific to MS and that might underlie its pathogenesis. Thus, the present study aims to provide an integrated evaluation of thymic function, peripheral T cell subsets homeostasis and of regulatory cells on treatment naïve RRMS patients at MS clinical onset.

## METHODS

Detailed methods available on eMethods

### Standard Protocol Approvals, Registrations, and Patient Consents

This project is in accordance with the Declaration of Helsinki and was approved by the local ethical committees of *Hospital de Braga* (ref. 5888/2016-CESHB), *Centro Hospitalar Universitário do Porto* (ref. 098-DEFI/097-CES) and *Hospital Álvaro Cunqueiro* (ref. 2021/430). All individuals that agreed to participate in the study signed a written informed consent.

### Data availability statement

The data that support the findings of this study are available from the corresponding author upon request.

## RESULTS

### Thymic export is unaltered in newly diagnosed RRMS patients

To evaluate if the thymic function, including intrathymic proliferation and thymic export, is altered in newly diagnosed RRMS patients, two surrogates were quantified in the blood: i) percentage and number of RTEs, defined as CD31^+^ naïve CD4^+^ T cells; and ii) levels of T cell receptor (TCR) excision circles (TRECs; episomal DNA molecules formed upon TCR locus rearrangement; absolute quantification of sjTREC is a measure of thymic export, whereas the sj/DJβTRECs ratio is a measure for intrathymic proliferation^28^). RTEs percentage and absolute numbers were unaltered in patients comparing to controls (**Figure 1A**). No differences were found on the sj/DJβTRECs ratio, nor on the sjTRECs and DJβTRECs levels (**Figure 1B-D; eTable 1, 2**). Overall, these results suggest that thymic function is not altered in newly diagnosed RRMS patients.

**Figure 1.**
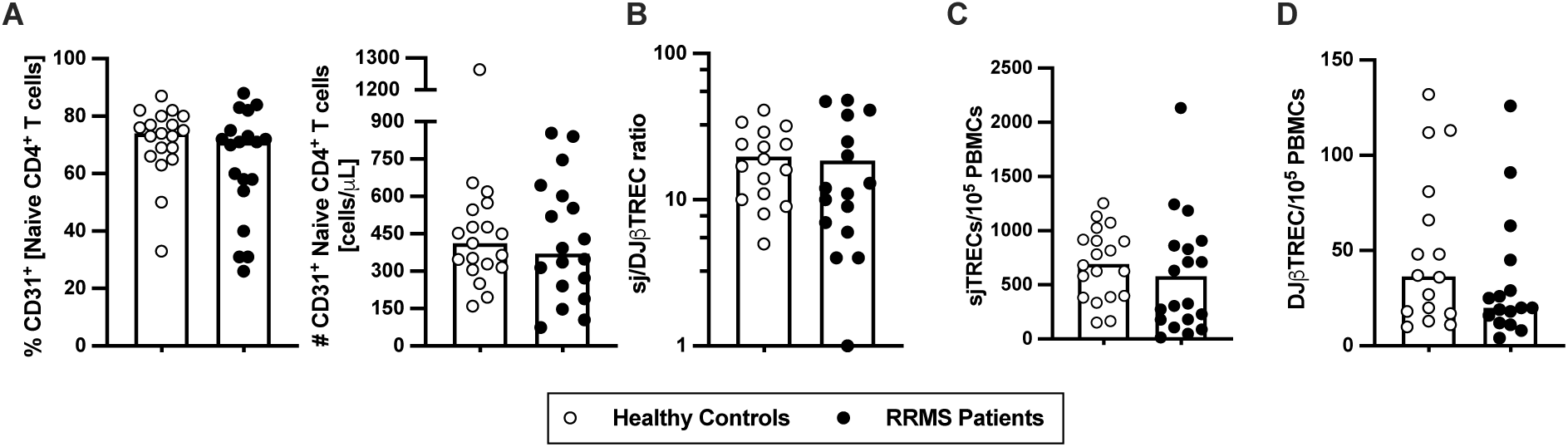
Thymic export in newly diagnosed RRMS patients does not differ from healthy controls. Percentage and number of CD31^+^ naïve CD4^+^ T cells (**A**). Ratio of sj/DJβTRECs (**B**) calculated from the quantification of sjTREC (**C**) and of DJβTREC (**D**) levels per 10^5^ peripheral blood mononuclear cells. In all graphs, each dot represents one individual, and the horizontal lines the group’s mean or median, as the data follow a normal or non-normal distribution, respectively. Comparisons between healthy controls (white circles) and newly diagnosed relapse-remitting multiple sclerosis (RRMS) patients (black circles) were performed using the t-test in **B** and the non-parametric Mann-Whitney *U*-test in **A, C** and **D** (statistical outputs and effect size calculations in eTable 1). When differences were statistically significant (*p-value* <0.050), the *p*-value was represented by * for 0.010< p <0.050; ** 0.001< *p* δ0.010; and *** *p* δ0.001. Differences are maintained upon controlling for sex, age, and human cytomegalovirus IgG seroprevalence on multiple linear regression models (eTable 2).

### Newly diagnosed RRMS patients have lower numbers of circulating memory T cells

As RTEs contribute to the peripheral seeding of the naïve T cell compartment and maintenance of naïve and memory T cells homeostasis^29^, the circulating levels of naïve and memory CD4^+^ and CD8^+^ T cells were evaluated. The percentage and number of total CD4^+^ T cells did not differ between newly diagnosed RRMS patients and HC (**Figure 2A; eTables 1, 3**). Based on CCR7 and CD45RA expression the naïve, central memory, effector memory and terminally differentiated CD45RA-expressing memory T cells (TemRA) were defined (**Figure 2B**). Among CD4^+^ T cells, no differences were found on the naïve/memory ratio (**Figure 2C**), neither on the percentages and numbers of naïve (**Figure 2D**), overall memory (**Figure 2E**) and central memory cells (**Figure 2F**). However, despite no differences are observe on the number and percentages of effector memory and TemRA CD4^+^ T cells upon direct comparison (**Figure 1G, 1F** and **eTable 1**), RRMS patients present lower numbers of effector memory CD4^+^ T cells (**eTable 3**), when data are adjusted for age, sex and HCMV IgG seroprevalence, using multiple linear regression models. Regarding the T helper (Th) phenotype, defined based on the differential expression of the chemokine receptors CCR4, CCR6, CCR10 and CxCR3,^5^ no differences were found on the percentages or numbers of Th1, Th2, Th9, Th17, Th22 and GM-CSF-producing Th cells (ThG) (**eFigure 1; eTable 1, 4**). No differences were found between groups regarding the percentage or number of the memory CD4^+^ T cells subsets expressing each of the chemokine receptors (data not shown).

**Figure 2.**
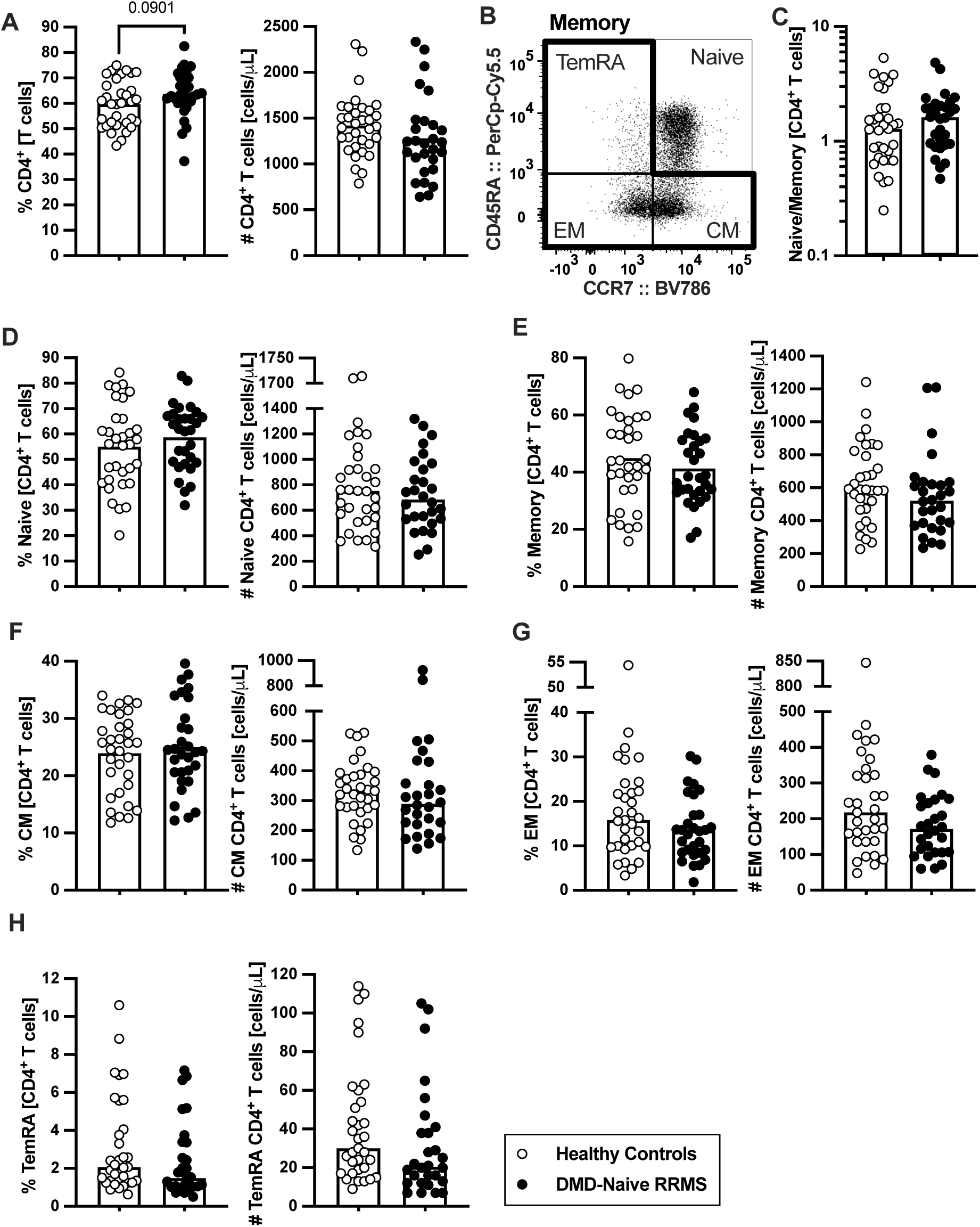
Newly diagnosed RRMS patients present no major differences on CD4^+^ T cells subsets. Percentages and numbers of CD4^+^ T cells are represented for healthy controls (white circles) and newly diagnosed relapse-remitting multiple sclerosis (RRMS) patients (black circles; **A**). Representative dot plot of the CD4^+^ T cell subpopulations (**B**): naïve (CD45RA^+^CD45RO^-^ CCR7^+^); central memory (CM; CD45RA^-^CD45RO^+^CCR7^+^), effector memory (EM; CD45RA^-^CD45RO^+^CCR7^-^); and terminally differentiated CD45RA-expressing memory T cells (TemRA; CD45RA^+^CD45RO^-^CCR7^-^). Total memory CD4^+^ T cells is the sum of the CM, EM and TemRA subpopulations. Naïve to memory ratio of CD4^+^ T cells (**C**). Percentages and numbers of CD4^+^ T cell subpopulations: naïve (**D**), total memory (**E**), central memory (**F**), effector memory (**G**) and TemRA (**H**). The parametric Student’s *t-*test was performed in **A, D** (percentage), **E** (percentage) and **F** (percentage); the non-parametric Mann-Whitney *U*-test in **C, D** (number), **E** (number), **F** (number), **G** and **H** (statistical outputs and effect size calculations in eTable 1). In all graphs, each dot represents one individual, and the horizontal lines the groups’ means or medians depending on the normal or non-normal distribution of the data, respectively. When differences were statistically significant (*p-value* <0.05), the *p*-value was represented by * for 0.010< p <0.050; ** 0.001< *p* ≤0.010; and *** *p* ≤0.001. Differences are maintained upon controlling for sex, age, and human cytomegalovirus (HCMV) IgG seroprevalence on multiple linear regression models (eTable 3), except for the percentage of CD4^+^ T cells (A), whose tendency to be higher in patients is lost and for the effector memory T cells whose absolute number becomes significantly lower in patients (G).

Lower number of CD8^+^ T cells was found in RRMS patients (**Figure 3A; eTable 1** and **5**). Concerning the naïve and memory subsets (**Figure 3B**), patients presented a tendency to lower naïve to memory CD8^+^ T cells’ ratio which was found to be associated with having the disease in the multiple linear regression model (**Figure 3C; eTable 1** and **5**). Though no differences were observed on the number of naïve CD8^+^ T cells (**Figure 3D; eTable 1** and **5**), RRMS patients present lower percentage and numbers of memory cells (**Figure 3E; eTable 1** and **5**). Regarding the memory CD8^+^ T cell subsets (**Figure 3B, F-H; eTable 1** and **5**), lower numbers of effector memory and TemRA cells were observed in patients (**Figure 3H** and **G**). We also found differences on CD8^+^ T cells expressing chemokine receptors, specifically the CCR6 (**eFigure 2A-D**). RRMS patients presented lower percentage and number of TemRA CD8^+^ T cells and lower number of effector memory CD8^+^ T cells expressing CCR6 (**eFigure 2A-D; eTable 1**), but the statistical significance of the latter was lost upon controlling for age, sex and HCMV IgG seroprevalence in the multiple linear regression model (**eTable 6**).

**Figure 3.**
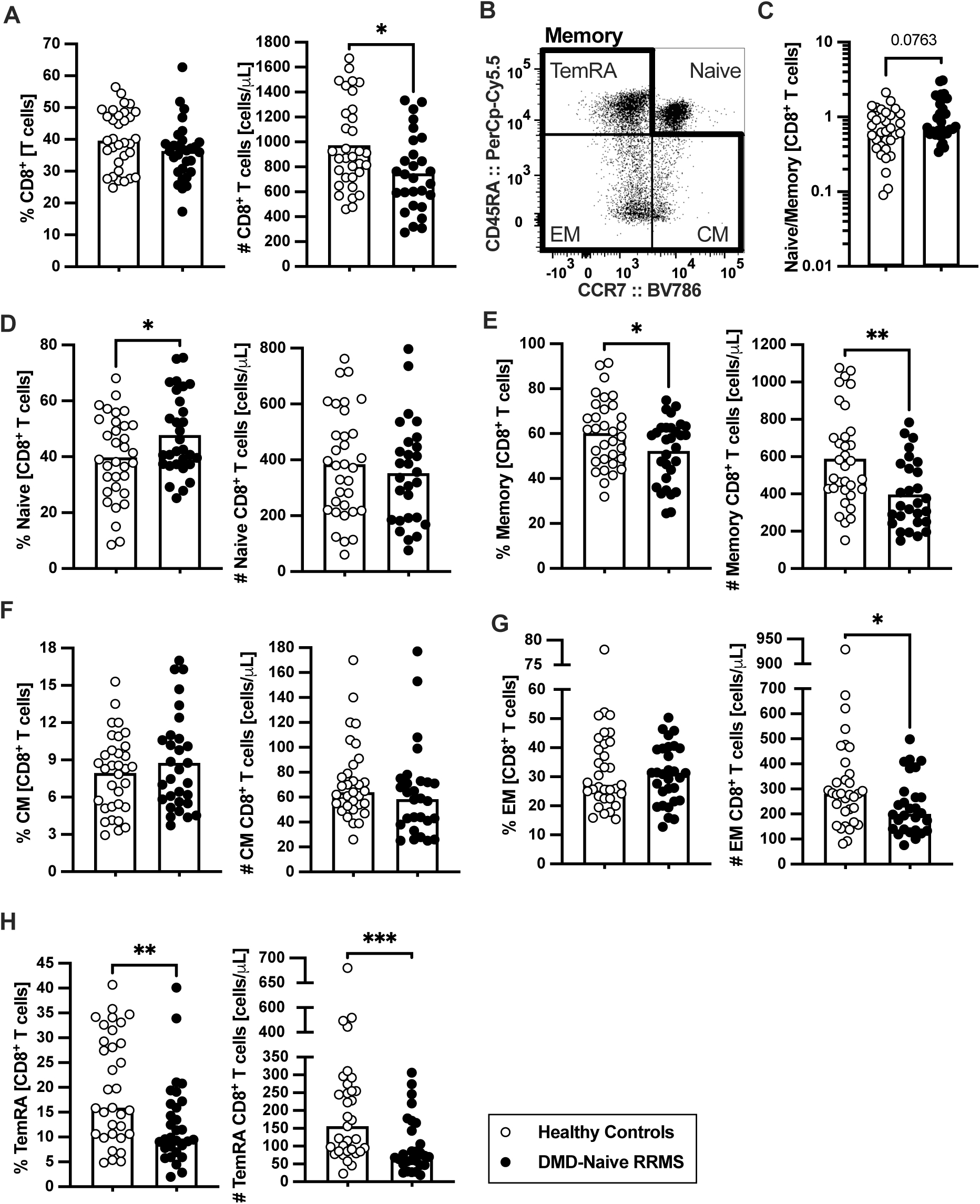
Newly diagnosed RRMS patients have lower numbers of memory CD8^+^ T cells and consequent higher naïve to memory ratio. Percentages and numbers of CD8^+^ T cells are represented for healthy controls (white circles) and newly diagnosed relapse-remitting multiple sclerosis (RRMS) patients (black circles; **A**). Representative dot plot of the CD8^+^ T cell subpopulations (**B**): naïve (CD45RA^+^CD45RO^-^CCR7^+^); central memory (CM; CD45RA^-^CD45RO^+^CCR7^+^), effector memory (EM; CD45RA^-^CD45RO^+^CCR7^-^); and terminally differentiated CD45RA-expressing memory T cells [TemRA; CD45RA^+^CD45RO^-^CCR7^-^). Total memory CD8^+^ T cells is the sum of the CM, EM and TemRA subpopulations. Naïve to memory ratio of CD8^+^ T cells (**C**). Percentages and numbers of CD8^+^ T cell subpopulations: naïve (**D**), total memory (**E**), central memory (**F**), effector memory (**G**) and TemRA (**H**). The parametric Student’s *t-*test was performed in **A** (number), **D, E, F** (percentage); the non-parametric Mann-Whitney *U*-test in **A** (percentage), **C, F** (number), **G** and **H** (statistical outputs and effect size calculations in eTable 1). In all graphs, each dot represents one individual, and the horizontal lines the groups’ means or medians depending on the normal or non-normal distribution of the data, respectively. When differences were statistically significant (*p-value* <0.050), the *p*-value was represented by * for 0.010< p <0.050; ** 0.001< *p* ≤0.010; and *** *p* ≤0.001. Differences are maintained upon controlling for sex, age, and human cytomegalovirus IgG seroprevalence on multiple linear regression models (eTable 5).

In summary, newly diagnosed RRMS patients present a general contraction of the peripheral memory T cell compartment, particularly pronounced in the CD8^+^ T cell subset.

### Newly diagnosed RRMS patients have higher percentages of naïve regulatory T cells

Tregs are scarcely characterized at MS clinical onset and their role on MS pathogenesis is yet unknown. In newly diagnosed RRMS patients no differences were found on the percentage or absolute number of Tregs comparing to HC (**Figure 4A; eTable 1, 7**). HLA-DR expression level is described to positively correlate with FOXP3 expression and with Tregs suppressive capacity.^30^ Three subsets of Tregs were defined based on HLA-DR and CD45RA expression: the naïve (CD45RA^+^HLA-DR^-^), and the activated HLA-DR^-^ (CD45RA^-^HLA-DR^-^) and HLA-DR^+^ (CD45RA^-^HLA-DR^+^) (**Figure 4B**). Higher naïve/activated ratio was observed in patients (**Figure 4C**). Regarding naïve Tregs, patients presented higher percentages of these cells, but no differences in numbers (**Figure 4D**). A tendency to lower percentage of activated HLA-DR^-^ Tregs was found in patients (**Figure 4E**; also observed in the regression models, **eTable 7**). No differences were found on activated HLA-DR^+^ Tregs (**Figure 4F**) though, upon controlling for age, sex and HCMV IgG seroprevalence, newly diagnosed RRMS patients exhibited a tendency to lower percentage of activated HLA-DR^+^ Tregs (**eTable 7**).

**Figure 4.**
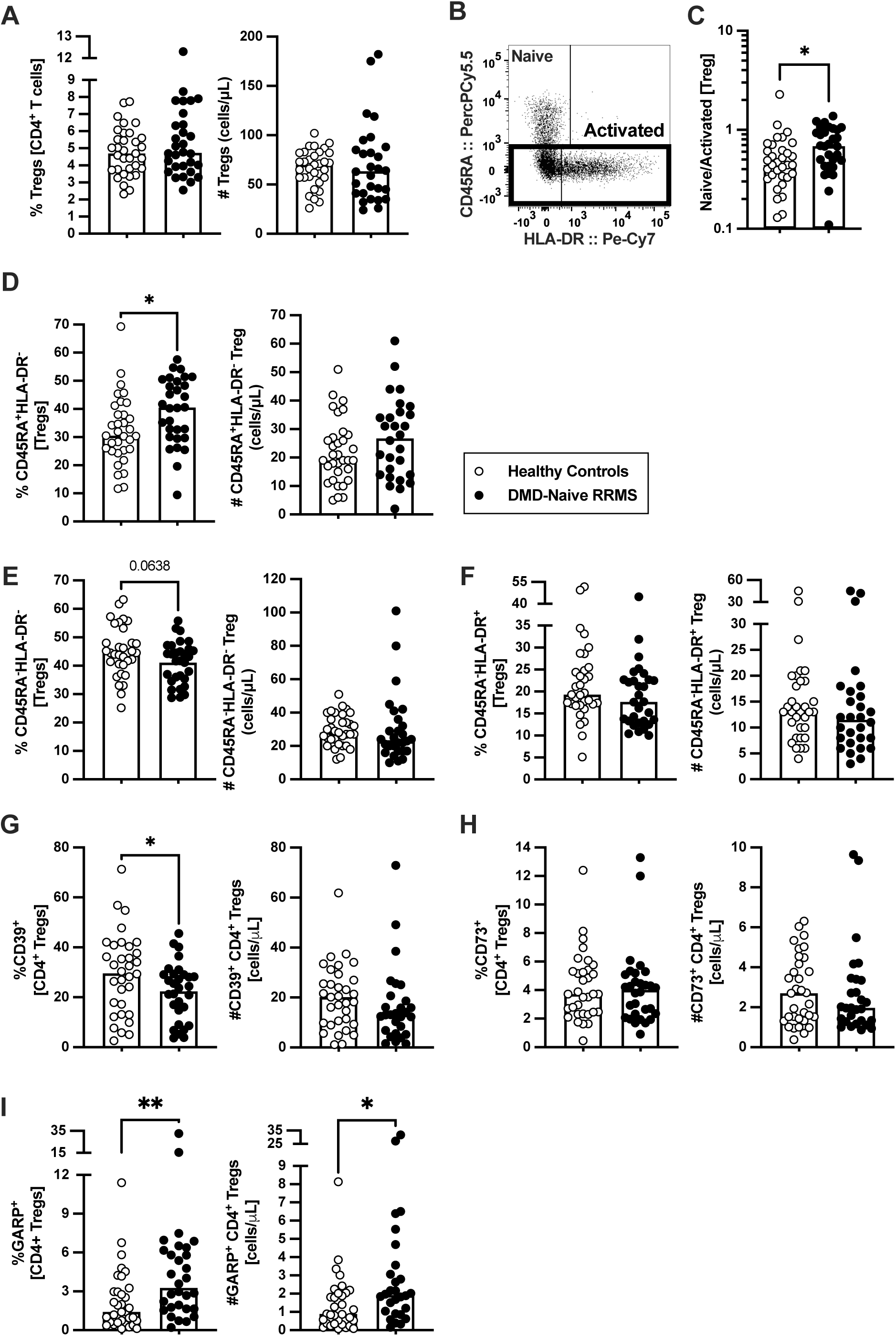
A higher naïve/activated Tregs’ ratio characterize newly diagnosed RRMS patients. Percentage and number of regulatory T cells (Tregs) are represented for healthy controls (white circles) and newly diagnosed relapse-remitting multiple sclerosis (RRMS) patients (black circles; **A**). Representative dot plot of Tregs subpopulations (**B**): naïve (CD45RA^+^HLA-DR^-^); activated HLA-DR^-^ (CD45RA^-^HLA-DR^-^); and activated HLA-DR^+^ (CD45RA^-^HLA-DR^+^). Total activated Tregs is the sum of the HLA-DR^-^ and HLA-DR^+^ subpopulations. Naïve/activated Tregs’ ratio (**C**). Percentages and numbers of Tregs subpopulations: naïve (**D**), activated HLA-DR^-^ (**E**), and activated HLA-DR^+^ (**F**). Percentages and numbers of Tregs expressing suppressive markers: CD39^+^ (**G**), CD73^+^ (**H**), and GARP^+^ (**I**).The parametric Student’s *t-*test was performed in **D** (number), **E** (percentage) and **G** (percentage); the non-parametric Mann-Whitney *U*-test in **A, C, D** (percentage), **E** (number),**F, G** (number), **H** and **I** (statistical outputs and effect size calculations in eTable 1). In all graphs, each dot represents one individual, and the horizontal lines the groups’ means or medians depending on the normal or non-normal distribution of the data, respectively. When differences were statistically significant (*p-value* <0.050), the *p*-value was represented by * for 0.010< p <0.050; ** 0.001< *p* ≤0.010; and *** *p* ≤0.001. Differences are maintained upon controlling for sex, age, and human cytomegalovirus (HCMV) IgG serology on multiple linear regression models (eTable 7), except for the naïve/activated Tregs’ ratio that becomes a tendency to be higher on RRMS patients (C), and the percentage of activated HLA-DR^+^ Tregs that becomes a tendency to be lower in patients (F).

The suppressive Treg markers CD39, CD73 and GARP were also evaluated to assess Tregs suppressive potential. Newly diagnosed RRMS patients presented lower percentage of CD39^+^ Tregs (**Figure 4G)** and no differences on CD73^+^ Tregs (**Figure 4H)**. A higher percentage and number of Tregs expressing GARP was observed in patients (**Figure 4I)**. Notwithstanding, the multiple linear regression model for the number of GARP-expressing Tregs was not statistically significant (**eTable 7)**.

These results point towards newly diagnosed RRMS patients having Tregs with a more naïve phenotype than HC.

### RRMS patients at clinical diagnosis have higher percentages of CD56^bright^ NK cells expressing KLRG1^+^ and of CD56^dim^CD57^+^ NK cells expressing NKp30^+^

NK cells characterization at clinical RRMS onset is unknown and poorly explored. At clinical diagnosis, RRMS patients were no different from HC on the percentage and number of NK cells (**Figure 5A**), or on any of its subsets [CD56^bright^ (most immature), CD56^dim^CD57^-^ and CD56^dim^CD57^+^ (most differentiated); **Figure 5B-D**)]. Newly diagnosed RRMS patients had higher percentage of KLRG1^+^ cells among CD56^bright^ NK cells (**Figure 5E**) but no differences on the percentage of cells expressing any of the other inhibitory receptors evaluated, such as NKG2A, KIR2DL2/3 and KIRDL1 (**Figure 5F-H; eTables 1 and 8**). Regarding the expression of the activating receptors, higher percentage of cells expressing NKp30^+^ cells was found among CD56^dim^CD57^+^ NK cells in patients (**Figure 5I**). No differences were observed on the percentage of cells expressing NKp44 or NKp46 receptors for any of the NK cell subsets (**Figure 5J-K; eTables 1 and 8**).

**Figure 5.**
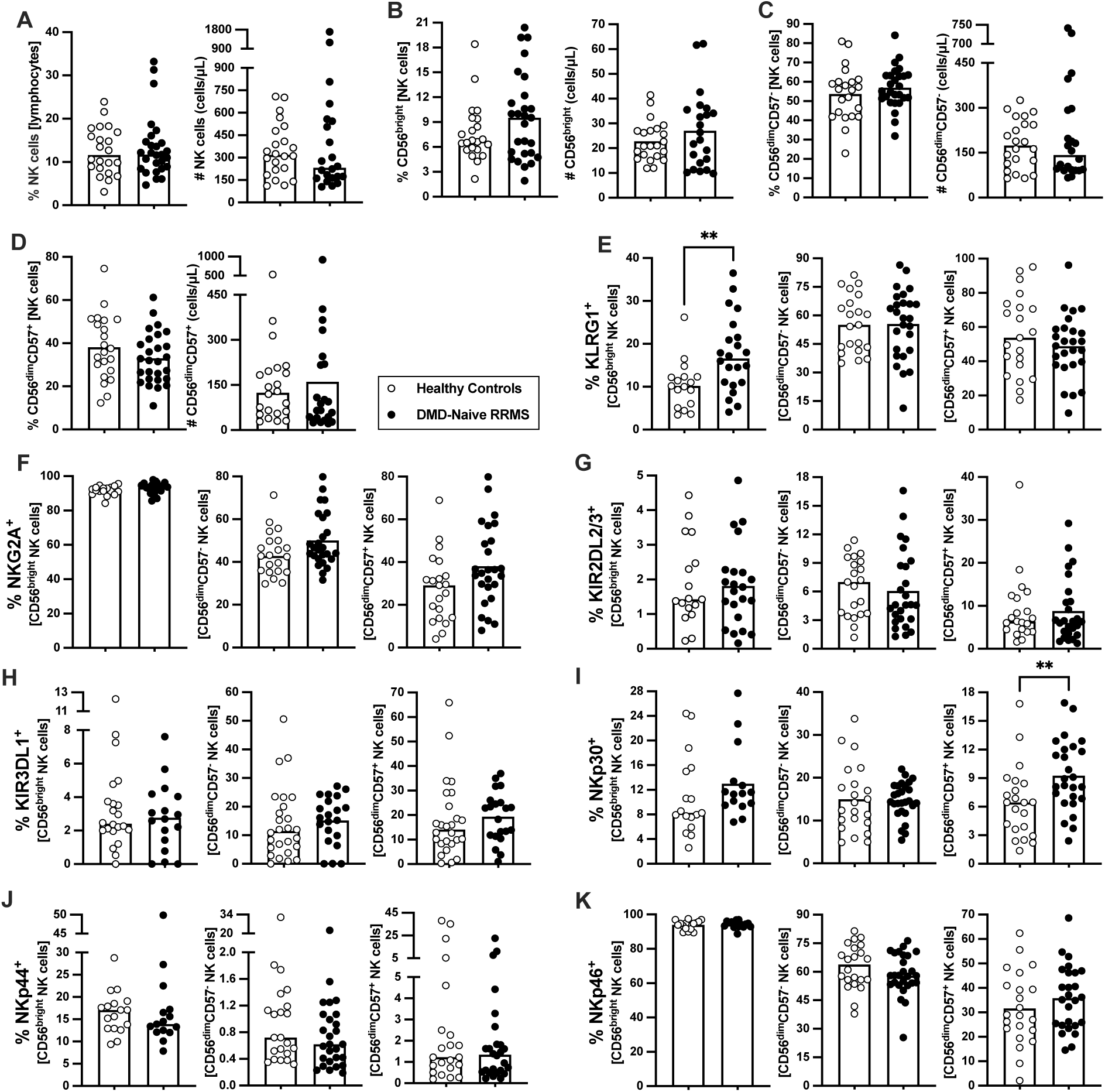
Newly diagnosed RRMS patients have higher percentages of CD56^bright^ NK cells expressing KLRG1 and CD56^dim^CD57^+^ expressing NKp30. Percentages and numbers of natural killer (NK) are represented for healthy controls (white circles) and newly diagnosed relapse-remitting multiple sclerosis (RRMS) patients (black circles; **A**). Percentages and numbers of NK cells subpopulations: CD56^bright^ (most immature; **B**); CD56^dim^CD57^-^ (**C**); and CD56^dim^CD57^+^ (most differentiated; **D**). Percentage of cells expressing inhibitory receptors in the three NK cells subsets: KLRG1 (**E**), NKG2A (**F**), KIR2DL2/3 (**G**) and KIR3DL1 (**H**). Percentage of cells expressing activating receptors in the three NK cells subsets: NKp30 (**I**), NKp44 (**J**) and NKp46 (**K**). The parametric Student’s *t-*test was performed in **B** (number), **C** (percentage), **D** (percentage), **E (**CD56^dim^CD57^-^ and CD56^dim^CD57^+^), **F** (CD56^dim^CD57^-^ and CD56^dim^CD57^+^), **G** (CD56^bright^ and CD56^dim^CD57^-^), **I** (CD56^dim^CD57^-^) and **K** (CD56^bright^ and CD56^dim^CD57^+^); the non-parametric Mann-Whitney *U*-test in **A, B** (percentage), **C** (number), **D** (number), **E** (CD56^bright^), **F** (CD56^bright^), **G** (CD56^dim^CD57^+^), **H, I** (CD56^bright^ and CD56^dim^CD57^+^), **J** and **K** (CD56^dim^CD57^-^) (statistical outputs and effect size calculations in eTable 1). In all graphs, each dot represents one individual, and the horizontal lines the groups’ means or medians depending on the normal or non-normal distribution of the data, respectively. When differences were statistically significant (*p-value* <0.050), the *p*-value was represented by * for 0.010< p <0.050; ** 0.001< *p* ≤0.010; and *** *p* ≤0.001. Differences are maintained upon controlling for sex, age, and human cytomegalovirus IgG serology on multiple linear regression models (eTable 8).

Newly diagnosed RRMS patients present no major differences on NKT cells concerning percentages and numbers (**eFigure 3A**). No differences were observed on percentage of NKT cells expressing of the inhibitory/activating receptors (**eFigure 3B-H**). The lower percentages of NKT cells expressing KIR3DL1 (**eFigure 3E)** was found to be related to HCMV IgG seroprevalence and not with the disease (**eTable 9**).

These results show that NK cells from newly diagnosed RRMS patients have increased percentages of the inhibitory KLRG1^+^ within the most immature NK cells subset and increased percentages of the activating NKp30^+^ within the most mature NK cells subset.

### Time from relapse and MS severity score (MSSS) correlate with alterations in immune cell populations

In RRMS, a relapse is a clinical manifestation of disease activity and results from an immune reaction that drives myelin destruction, and consequent neuronal damage and impairment.^1^ All patients from this cohort were on remission at the time of the immune characterization, though the vast majority (78%; 21 patients) had a relapse within the 10 months prior sampling. We next analyzed whether the time from the last relapse was correlated with peripheral immune alterations. As some patients had a corticoid pulse to treat the relapse, this was included as a dichotomic variable in the regression models evaluating the contribution of the time from last relapse to immune cells alterations to control for this possible confounding effect (**eTable 10**). Alterations on CD4^+^ and CD8^+^ T cells, Tregs, NK cells and NKT cell subsets were found to be related with time from the relapse (**Figure 6A; eTable 10**). The increasing time from relapse was associated with higher numbers of total CD4^+^ T cells, both naïve and total memory (specifically central memory and Th2 cells). Growing numbers of total memory CD4^+^ T cells expressing CCR4, and of CCR4^+^ and CCR10^+^ central memory cells were also allied with increasing time from relapse. Lower percentage of effector memory CD4^+^ T cells expressing CCR6^+^ was associated with increasing time from relapse. Regarding CD8^+^ T cells, higher percentage and number of central memory cells was related with increasing time from relapse. Also, increasing time from relapse was correlated with increased numbers and percentages of CCR10^+^ and CCR4^+^ cells among total, central and effector memory CD8^+^ T cells, and with a decrease in the percentages of CCR6^+^ cells in those cell subsets.

**Figure 6.**
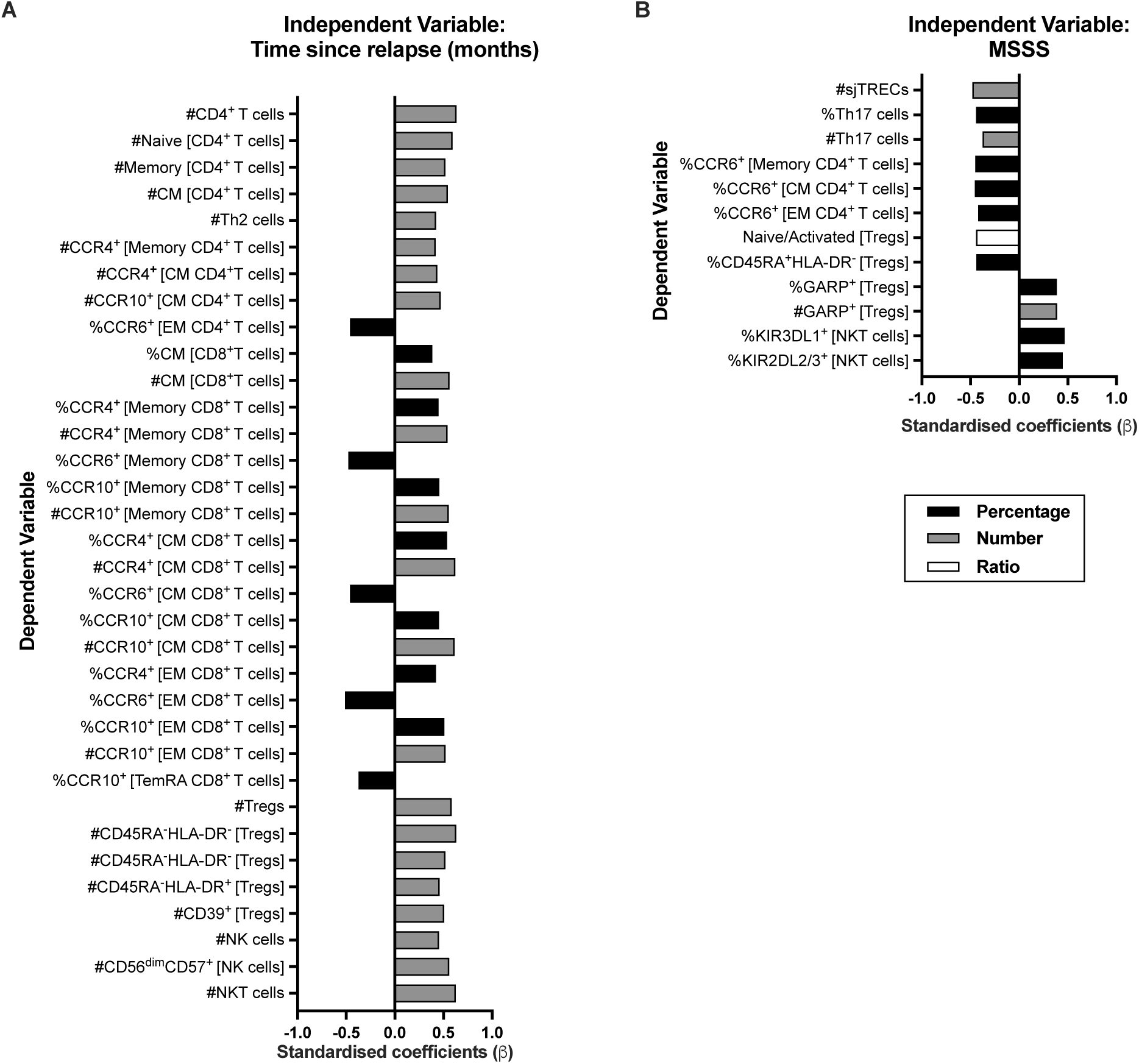
Time from the last relapse and MS severity score (MSSS) relates to immune system cell alterations. Represented are the standardized coefficients (β) from multiple or linear regression models having as an independent variable the time from last relapse (**A**) or the MSSS (**B**), respectively. The dependent variable consisted of the unstandardized residuals from a preliminary multiple linear regression where the contribution of sex, age and HCMV IgG seroprevalence on the blood populations was evaluated (as described in the eMethods section). Only the blood populations for which the linear regression models were significant (*p* <0.050) for the variable of interest are represented. Statistical outputs and effect size calculations are represented in eTables 10, 11.

Increasing numbers of total Tregs, of all its subsets based on HLA-DR and CD45RA, and of Tregs expressing CD39 were associated with increasing time from relapse. Higher numbers of NKT cells, NK cells and of CD56^dim^CD57^+^ (the most differentiated NK subset) correlated with increasing time from last relapse. We also investigated a possible association between the MSSS and the alterations on blood cell populations (**Figure 6B; eTable 11**). A higher MSSS correlated with lower sjTRECs levels and percentages of CCR6^+^ cells among memory CD4^+^ T cells (specifically central and effector memory), including, lower percentages and number of Th17 cells. Lower naïve/activated Tregs’ ratio and percentage of naïve Tregs (CD45RA^+^HLA-DR^-^) correlated with increasing MSSS. Higher percentage and number of Tregs expressing GARP was allied to higher MSSS. On NKT cells, higher MSSS correlated with higher percentage of cells expressing KIR3DL1 and KIR2DL2/3 inhibitory receptors. No association was found between MS severity and CD8^+^ T cells subsets.

## Discussion

Several reports have evaluated thymic function and blood cell populations in MS patients.^7–15^ Most of those studies described an aged immune system in MS patients in comparison to HC as assessed by thymic function, naïve to memory T cells’ ratio, among other parameters. However, in some of those studies, the group of MS patients is quite heterogeneous, combining in the same set patients with different MS progression forms^7,15^, diverse disease duration^10^ or RRMS patients on relapse with patients on remission.^11^ Additionally, in some instances, information regarding the MS treatment history is missing which hampers our comprehension on whether the effect of immunotherapies on immune cell populations was considered or neglected.^15^ Here we based our study on a homogenous group of newly diagnosed RRMS patients (<17 months [median of 1 month] after disease diagnosis), naïve for disease modifying drugs and recruited from three distinct hospitals. The study of these newly diagnosed RRMS patients, and of age and sex-matched HC allowed us to provide an integrative view of thymic function and of the several blood immune cell populations at MS clinical onset. Additionally, it adds on the relation between immune alterations and time from last relapse and MS severity. All these observations were controlled for age, sex and HCMV IgG seroprevalence, which are variables known to affect the percentage and numbers of blood cell populations.^20,31^

We report here similar thymic function on newly diagnosed RRMS patients and HC upon controlling for age, sex and HCMV IgG seroprevalence, as observed by equivalent levels of RTEs, sjTRECs, DJβTRECs and sj/DJβTREC ratio. Despite many studies support a lower thymic function in RRMS patients,^8,11–13^ at least one other report corroborates our data on untreated RRMS patients.^14^ Moreover, in a cohort of treatment-naïve monozygotic twins discordant for MS it was found that RRMS itself was not associated with RTEs alterations.^32^ We found that higher MSSS are related to lower sjTRECs levels in patients, but not with the sjTREC/DJβTREC ratio. Unlike sjTREC/DJβTREC ratio, sjTRECs quantification, by itself, is not a good surrogate of thymic function as cells containing sjTRECs get diluted with cell proliferation.^28^ For this reason, the alteration on sjTRECs with MSSS should not be related with altered thymic function.

We observed that newly diagnosed RRMS patients present lower number of effector memory CD4^+^ T cells upon controlling for sex, age and HCMV seroprevalence. A previous study reported that untreated RRMS patients had lower percentage, and a tendency to lower number, of early effector memory CD4^+^ T cells and speculated that those alterations were attributed to T cell migration into the CNS.^14^ Curiously, another study observed a cluster of central/effector memory T cells with a CNS homing signature that was reduced in the blood of MS patients, and enriched in lesions of post-mortem brain tissue of MS patients, in comparison to HC.^33^ The magnitude of the observed differences suggest a systemic alteration on CD4^+^ T cells, and not specifically on myelin specific CD4^+^ T cells since these cells represent a negligible population of approximately 0.0001% of all CD4^+^ T cells and the percentage of autoreactive CD4^+^ T cells in blood has been described to not differ between RRMS and healthy individuals.^34^ From our data we cannot infer on whether the lower number of effector memory T cells is related to altered proneness to migrate to tissues as no major differences were found between groups on the expression of chemokine receptors, nor on any of the other memory CD4^+^ T cell subsets (data not shown). Yet, fewer effector memory CD4^+^ T cells expressing CCR6 were found with increasing time from relapse. As CCR6 has been pointed as a chemokine receptor involved in the migration of cells to the CNS, those findings might suggest that during a relapse there are more effector memory CD4^+^ T cells with potential to migrate to the CNS.^35^ As time from the last relapse increases, higher numbers of blood CD4^+^ T cells were also observed, particularly of the naïve and central memory cells, pointing towards a possible involvement of these cells on MS relapses, though the link between naïve and central memory T cells reduction and MS pathophysiology was not explored.

Th1 and Th17 cells have been suggested to be involved in MS pathophysiology.^4^ Newly diagnosed RRMS patients did not differ from HC on the percentage or number of Th1 and Th17, nor on the balance of Th17/Tregs (data not shown), whose imbalance has been implicated in autoimmunity.^36^ Still, we observed a correlation between lower percentage and number of Th17 cells and higher MSSS; whether this is due to migration and accumulation of those cells on the CNS of patients and whether that impacts disease severity must be further explored.

We observed lower number of CD8^+^ T cells in the blood of RRMS patients. CD8^+^ T cells are known to surpass CD4^+^ T cells in MS lesions, which may imply a higher recruitment of CD8^+^ T cells to the CNS.^6^ Our data show decreased numbers of CCR6-expressing memory CD8^+^ T cells in patients, which might be related to increased migration of these cells to tissues. Interestingly, CCR6 has been described to facilitate brain-homing migration and to be essential for EAE induction.^35^ Additionally, the choroid plexus constitutively expresses the CCR6-ligand CCL20 and constitutes the most probable route of entry for those cells into the CNS.^35^ Interestingly, lower percentages of memory CD8^+^ T cells expressing CCR6 were associated with increasing time from relapse, which suggests that during a relapse a higher percentage of cells bearing brain-homing potential might be present in circulation. We observed that RRMS patients present higher naïve/memory ratio among CD8^+^ T cells and lower number of memory T cells, specifically of effector memory and TemRA. Lower percentage of these memory CD8^+^ T cells subsets was previously observed in a group of untreated RRMS and Clinically Isolated Syndrome (CIS) patients, in comparison to healthy controls.^37^ The authors attributed the lower effector memory and TemRA CD8^+^ T cells to primary intrinsic defects (genetically determined) rather than being a pathological consequence of MS. Additionally, the authors claim that the memory CD8^+^ T cell deficiency is present since MS onset and persists throughout its course independently of disease severity, disability, or duration.^37^ Curiously, CD8^+^ T cells deficiency is characteristic of many other chronic autoimmune disorders (e.g. rheumatoid arthritis, systemic lupus erythematosus, Crohn’s disease, type 1 diabetes mellitus, myasthenia gravis, etc.) and is also found in the blood of patients’ relatives.^38^ This supports that CD8^+^ T cells deficiency might be inherited and underly autoimmunity.

The essential role of Tregs to immune tolerance highlights the importance of these cells in autoimmune disorders.^17^ Our data show that in comparison to HC, newly diagnosed RRMS patients present higher percentage of naïve Tregs, and tendency to lower percentages of HLA-DR^+^ and HLA-DR^-^ activated Tregs subsets. These results contrast with previous findings showing higher percentage of activated HLA-DR^+^ and lower of naïve Tregs in MS patients.^19,39^ Our focus specifically on newly diagnosed RRMS patients and the combination of CD25, CD127, and FOXP3 expression to define Tregs (and its subsets using HLA-DR and CD45RA) might have directed us to these observations. The percentage of cells expressing the suppressive marker CD39 was lower in patients’ Tregs, which might be related to the fact that this marker was mostly expressed on activated Tregs, that were also tendentially lower in RRMS patients. In fact, the percentage of CD39^+^ among activated Tregs was not different between groups (data not shown), thus suggesting that the suppressive capacity of activated Tregs is not altered, as assessed by CD39. On the other hand, the percentage and number of GARP-expressing Tregs is higher in RRMS patients, which is suggestive of higher proportion of suppressive Tregs in patients. Similar findings were previously made in RRMS patients, and the authors attributed it to systemic inflammation rather than being a functional defect of patients’ Tregs.^40^ A higher number of Tregs (and of all its subsets based on CD45RA and HLA-DR expression), and of Tregs expressing CD39 was correlated with increasing time from relapse. On the other hand, increasing MSSS were linked with decreased naïve/activated ratio and percentage of naïve Tregs, and with higher numbers of suppressive GARP^+^ Tregs.

Altogether, these results suggest that during a relapse fewer Tregs, including Tregs bearing suppressive potential through CD39 expression, are present in the blood of patients and that patients with higher MSSS have more Tregs expressing GARP and lower percentage of naïve Tregs. Though, a longitudinal study evaluating these cell subsets in naïve RRMS on relapse and in remission would be valuable to clarify these aspects. Notwithstanding, our results correlate with previous data showing that the activated CD45RA^-^FOXP3^hi^ Tregs increase with time from relapse,^39^ and that RRMS patients with lower percentages of naïve Tregs are the ones with highest expanded disability expanded score (EDSS), an alternative scale of disability to MSSS.^41^

The CD56^bright^ cells belong to the most immature NK cell subset, and are described to be more potent at producing cytokines.^42^ We observed that RRMS patients had higher percentage of these cells expressing the inhibitory receptor KLRG1. KLRG1 binds E-, N- and R-cadherins, being the last 2 found in the CNS. Specifically, N-cadherin was found to be increased in remyelinating lesions of EAE mice, particularly in oligodendrocytes.^43^ The increase in the percentage of KLRG1^+^ cells might suggest that the most immature NK subset, the CD56^bright^ NK cells, from patients might be less prone to kill target cells than the ones from healthy individuals, though functional studies should clarify this issue. Regarding the activation receptors, we observed that patients present a higher percentage of CD56^dim^CD57^+^ NK cells, most mature and cytotoxic NK cell subset, expressing NKp30. Engagement of the natural cytotoxicity receptors NKp30, NKp44 and/or NKp46 from CD56^bright^ NK cells is required to suppress autologous T cells from untreated RRMS patients *in vitro*.^23^ Yet, no significant suppression of autologous T cells from untreated RRMS patients was observed by the most mature CD56^dim^ NK cells.^23^ Even though, we cannot exclude a possible role of CD56^dim^CD57^+^ NK cells at controlling autoreactive T cells as a higher number of this cell subset was related with increasing time from relapse. Regarding NKT cells, despite RRMS patients had lower percentage of these cells expressing KIR3DL1, this difference was lost and mostly attributed to HCMV IgG seroprevalence. Still, higher numbers of total NKT cells were related with higher time from relapse, and higher percentages of these cells expressing the inhibitory receptors KIR2DL2/3 and KIR3DL1 are related to higher MS severity, an observation not previously described to our knowledge.

In summary, we observed profound peripheral immune alterations in newly diagnosed RRMS patients including unbalanced naïve to memory cells due to deficiency on the memory subset, particularly on CD8^+^ T cells. Furthermore, these patients were characterized by alterations on regulatory cells, presenting more naïve Tregs, higher percentages of immature NK cells expressing inhibitory receptors and higher percentages of the most mature NK cells expressing activating receptors. Altogether, an unbalance on conventional T cells together with unbalanced regulatory cells subsets seem to characterize RRMS patients at clinical diagnosis. Even though MS initiates much earlier than symptoms onset, the characterization of the peripheral immune system cells in newly diagnosed RRMS patients unveiled immune features present at disease clinical onset that might set the basis to the better understanding of the disease pathogenesis and the search for novel diagnostic biomarkers.

## FIGURE LEGENDS

**eFigure 1. Newly diagnosed RRMS patients present no differences on T helper (Th) subsets**. Percentages and numbers of Th subsets are represented for healthy controls (white circles) and newly diagnosed relapse-remitting multiple sclerosis (RRMS) patients (black circles). Percentages and numbers of Th cell subsets: Th1 (**A**), Th2 (**B**), Th9 (**C**), Th17 (**D**), Th22 (**E**) and Th-GM-CSF (ThG; **F**). The parametric Student’s *t-*test was performed in **B** (percentage) and **D** (percentage); the non-parametric Mann-Whitney *U*-test in **A, B** (number), **C, D** (number), **E** and **F** (statistical outputs and effect size calculations in eTable 1). In all graphs, each dot represents one individual, and the horizontal lines the groups’ means or medians depending on the normal or non-normal distribution of the data, respectively. When differences were statistically significant (*p-value* <0.05), the *p*-value was represented by * for 0.010< p <0.050; ** 0.001< *p* ≤0.010; and *** *p* ≤0.001. Differences are maintained upon controlling for sex, age, and human cytomegalovirus (HCMV) IgG seroprevalence on multiple linear regression models (eTable 4), except for the Th2 cells (B) whose number becomes tendentially lower in in patients.

**eFigure 2. Newly diagnosed RRMS patients present lower memory CD8**^**+**^ **T cells expressing the chemokine receptor CCR6**. Percentages and numbers of chemokine receptors expressing CD8^+^ T cells subsets are represented for healthy controls (white circles) and newly diagnosed relapse-remitting multiple sclerosis (RRMS) patients (black circles). Percentages and numbers of CCR4, CCR6, CCR10 and CxCR3 are represented among: total memory (**A**), central memory (**B**), effector memory (**C**) and TemRA (**D**). The parametric Student’s *t-*test was performed in **B** (CCR6 and CxCR3 percentages) and **D** (CxCR3 percentage); the non-parametric Mann-Whitney *U*-test in **A, B** (CCR6 and CxCR3 number and in CCR4 and CCR10), **C** and **D** (CCR6 number and in CCR4, CCR10 and CxCR3); statistical outputs and effect size calculations in eTable 1. In all graphs, each dot represents one individual, and the horizontal lines the groups’ means or medians depending on the normal or non-normal distribution of the data, respectively. When differences were statistically significant (*p-value* <0.05), the *p*-value was represented by * for 0.010< p <0.050; ** 0.001< *p* ≤0.010; and *** *p* ≤0.001. Differences are maintained upon controlling for sex, age, and human cytomegalovirus (HCMV) IgG seroprevalence on multiple linear regression models (eTable 6), except for the central memory CD8^+^ T cells expressing CCR6 (A), where a statistical tendency to lower number of those cells became evident in patients and for the effector memory CD8^+^ T cells expressing CCR6 (C), whose lower number observed in patients was lost due to sex and HCMV IgG seroprevalence.

**eFigure 3. Newly diagnosed RRMS patients have lower percentages of NKT cells expressing the inhibitory receptor KIR3DL1**. Percentages and numbers of natural killer (NK)T cells are represented for healthy controls (white circles) and newly diagnosed relapse-remitting multiple sclerosis (RRMS) patients (black circles; **A**). Percentage of cells expressing inhibitory receptors in NKT cells: KLRG1 (**B**), NKG2A (**C**), KIR2DL2/3 (**D**) and KIR3DL1 (**E**). Percentage of cells expressing activating receptors in NKT cells: NKp30 (**F**), NKp44 (**G**) and NKp46 (**H**). In all graphs, each dot represents one individual, and the horizontal lines the groups’ medians, as data follow a non-normal distribution. The non-parametric Mann-Whitney *U*-test was performed to compare groups (statistical outputs and effect size calculations in eTable 1). When differences were statistically significant (*p-value* <0.050), the *p*-value was represented by * for 0.010< p <0.050; ** 0.001< *p* δ0.010; and *** *p*δ0.001. Differences are maintained upon controlling for sex, age, and human cytomegalovirus (HCMV) IgG serology on multiple linear regression models (eTable 9), except for the percentage of KIR3DL1^+^ NKT cells where the significant difference is lost because of HCMV IgG serology (E).

**eFigure 4. Gating strategy for phenotypical analysis of recent thymic emigrants**. Issues during acquisition were verified by accompanying acquisition throughout time **(A)**. Leukocytes were then selected by their positive expression of CD45 (**B**), and viable cells selected by excluding 7AAD positive cells (**C**). After isolating the singlets (**D**), the lymphocytes were gated regarding their lower size (FSC-A) and complexity (SSC-A; **E**). Within the lymphocytes population T cells were defined through the expression of CD3^+^ (**F**). CD4^+^ were then discriminated among T cells (**G**). Gates for CD45RA and CCR7 were combined to discriminate the Naïve subset (CD45RA^+^CD45RO^-^CCR7^+^; **H**). Finally, recent thymic emigrants were defined by the expression of the CD31 marker among the naïve CD4^+^ T cell subset (CD45RA^+^CCR7^+^CD31^+^; **I**).

**eFigure 5. Gating strategy for phenotypical analysis of the main T cell subsets**. Issues during acquisition were verified and excluded using the FlowAI plugin in FlowJo. Singlets were isolated (**A**) and live leukocytes selected by their positive expression of CD45 and exclusion of the positive cells for the viability dye (**B**). The lymphocytes were gated regarding their lower size (FSC-A) and complexity (SSC-A; **C**). Within the lymphocyte population, T cells were defined by the positive expression of CD3 (**D**). CD4^+^ and CD8^+^ cell subsets were then discriminated among T cells (**E**). CD45RA and CCR7 were used to distinguish both for CD4^+^ (**H**) and CD8^+^ (**I**) T cells the Naïve (CD45RA^+^CCR7^+^), Central Memory (CD45RA^-^CCR7^+^), Effector Memory (CD45RA^-^CCR7^-^) and Effector Memory expressing CD45RA (TemRA; CD45RA^+^CCR7^-^) subsets [**F; G**]. Finally, T helper (Th) subsets were defined among total memory CD4^+^ T cells through the differential expression of CCR4, CCR6, CCR10 and CxCR3 (**H-K**): Th1 (CxCR3^+^); Th2 (CCR4^+^CCR6^-^CCR10^−^CxCR3^-^); Th9 (CCR6^+^CCR4^-^); Th17 (CCR4^+^CCR6^+^CCR10^−^CxCR3^-^); Th22 (CCR4^+^CCR6^+^CCR10^+^CxCR3^-^) and ThG (CCR4^+^CCR6^-^CCR10^+^CxCR3^-^).

**eFigure 6. Gating strategy for phenotypical analysis of regulatory T cells (Tregs)**. Issues during acquisition were verified and excluded using the FlowAI plugin in FlowJo. Singlets were selected (**A**) and live leukocytes discriminated by their positive expression of CD45 and by excluding the positive cells for the viability dye (**B**). The lymphocytes were gated according to their lower size (FSC-A) and complexity (SSC-A; **C**). Within the lymphocytes, T cells were defined by the expression of CD3^+^ (**D**) followed by discrimination of the CD4^+^ and CD8^+^ T cell subsets (**E**). Tregs were defined after selection of cells expressing lower levels of CD127 (**F**) followed by selection of FoxP3 and CD25 positive CD4^+^ T cells (**G**). The CD45RA and HLA-DR markers were used to define 3 Tregs subsets: the naive (CD45RA^+^HLA-DR^-^), the activated HLA-DR^-^ (CD45RA^-^HLA-DR^-^) and the activated HLA-DR^+^ (CD45RA^-^HLA-DR^+^) Tregs (**H**). Suppressive markers were also evaluated among total Tregs: CD39 (**I**), CD73 (**J**) and GARP (**K**).

**eFigure 7. Gating strategy for phenotypical analysis of NKT and NK subsets regarding expression of inhibitory and activating receptors**. Issues during acquisition were verified by accompanying acquisition throughout time **(A)**. Live leukocytes were then selected by their positive expression of CD45 and by excluding positive cells for the viability dye (**B**). After isolating the singlets (**C**), the lymphocytes were gated regarding their size (FSC-A) and complexity (SSC-A; **D**). Within the lymphocytes population NK and NKT cells were defined by their positive expression of CD56 and differential expression of CD3 (**E**). NK cell subsets were then discriminated according to the expression of CD57 and level of expression of CD56 into 3 main populations: CD56^bright^ NK cells, CD56^dim^CD57^-^ NK cells and CD56^dim^CD57^+^ NK cells. Depending on the staining panel the expression of inhibitory (KIR2DL2/3, KIR3DL1, KLRG1 and NKG2A) or of activating receptors (NKp30, NKp44 and NKp46) were evaluated within the NK subsets and NKT cells (**G**).

