## Supplementary figures and images for "Low memory T cells blood counts and high naïve regulatory T cells percentage at relapsing remitting multiple sclerosis diagnosis"

### Supplemental Figures

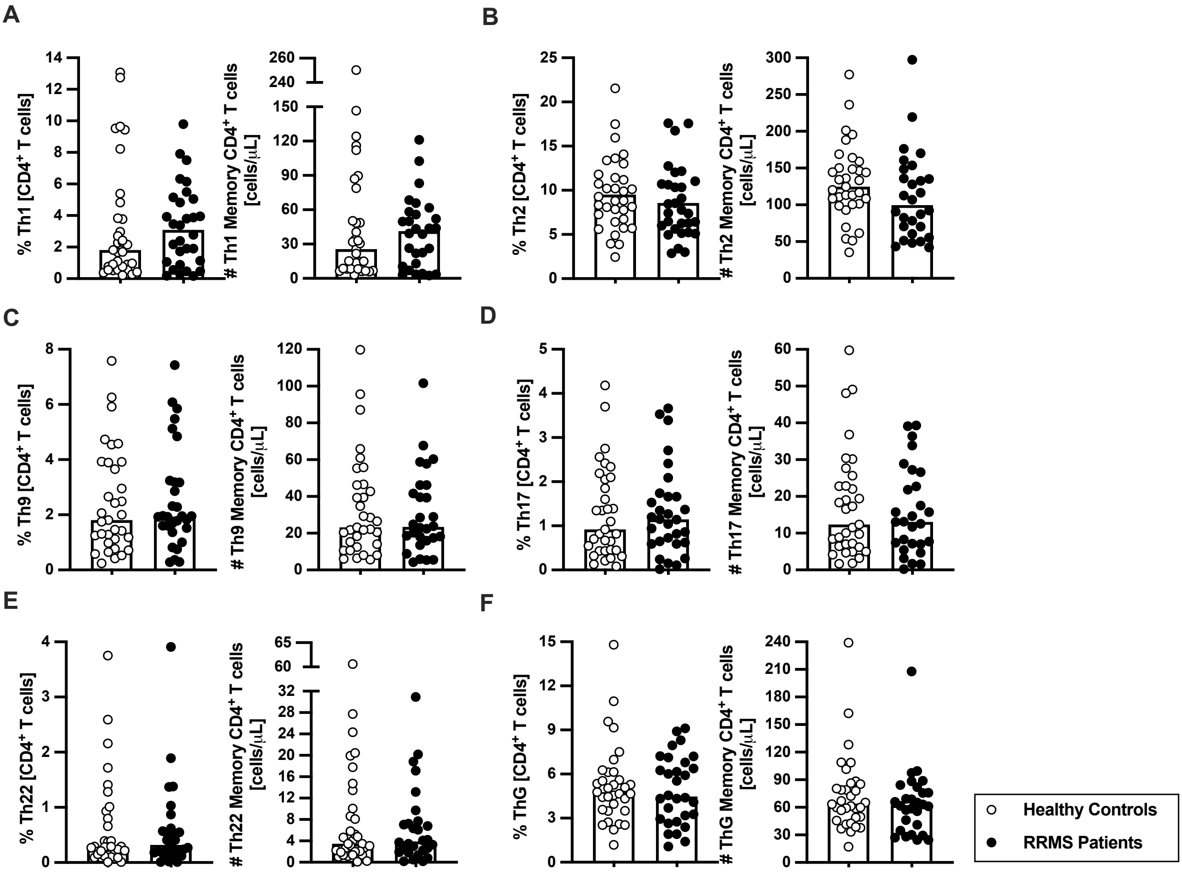

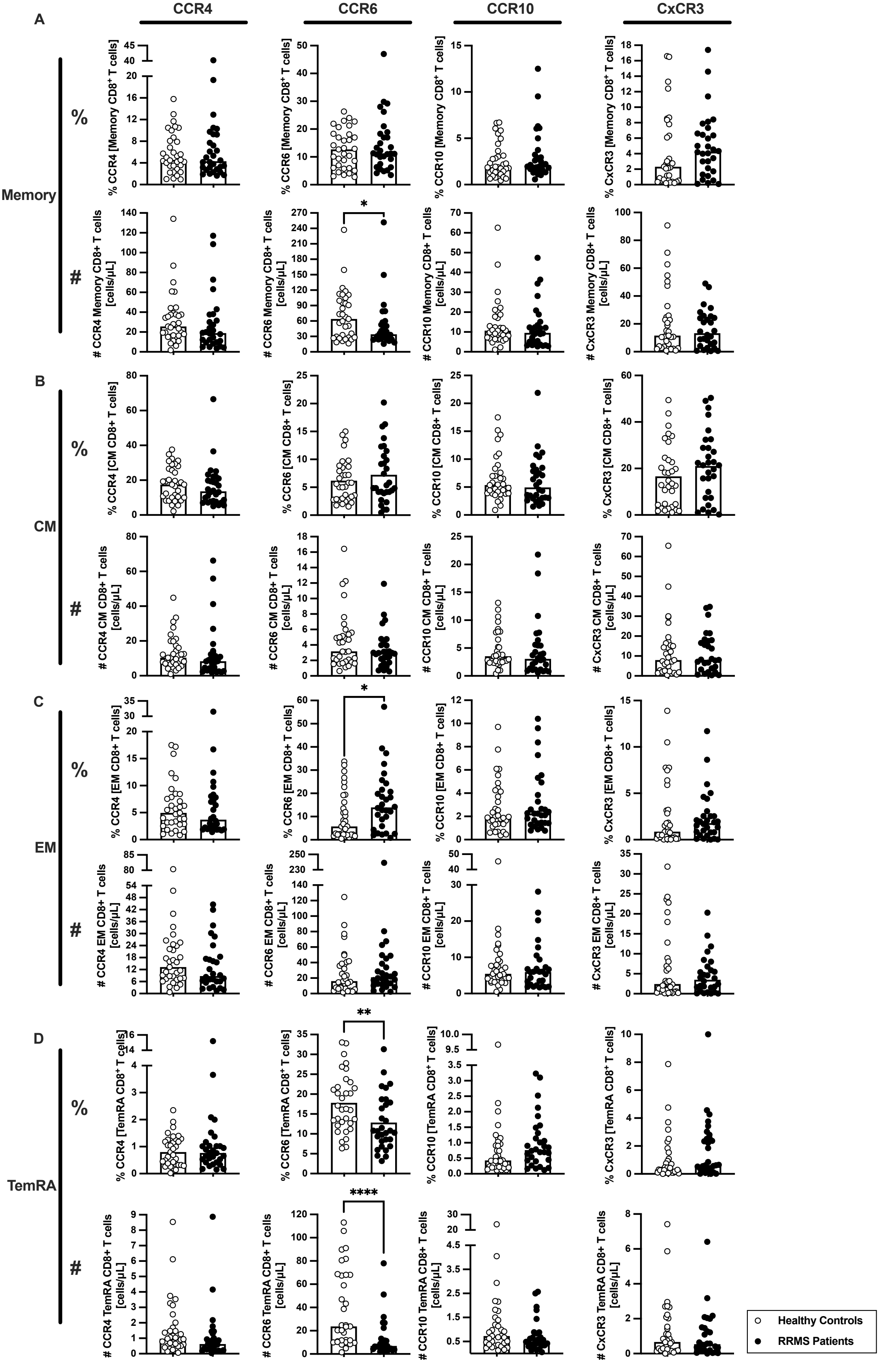

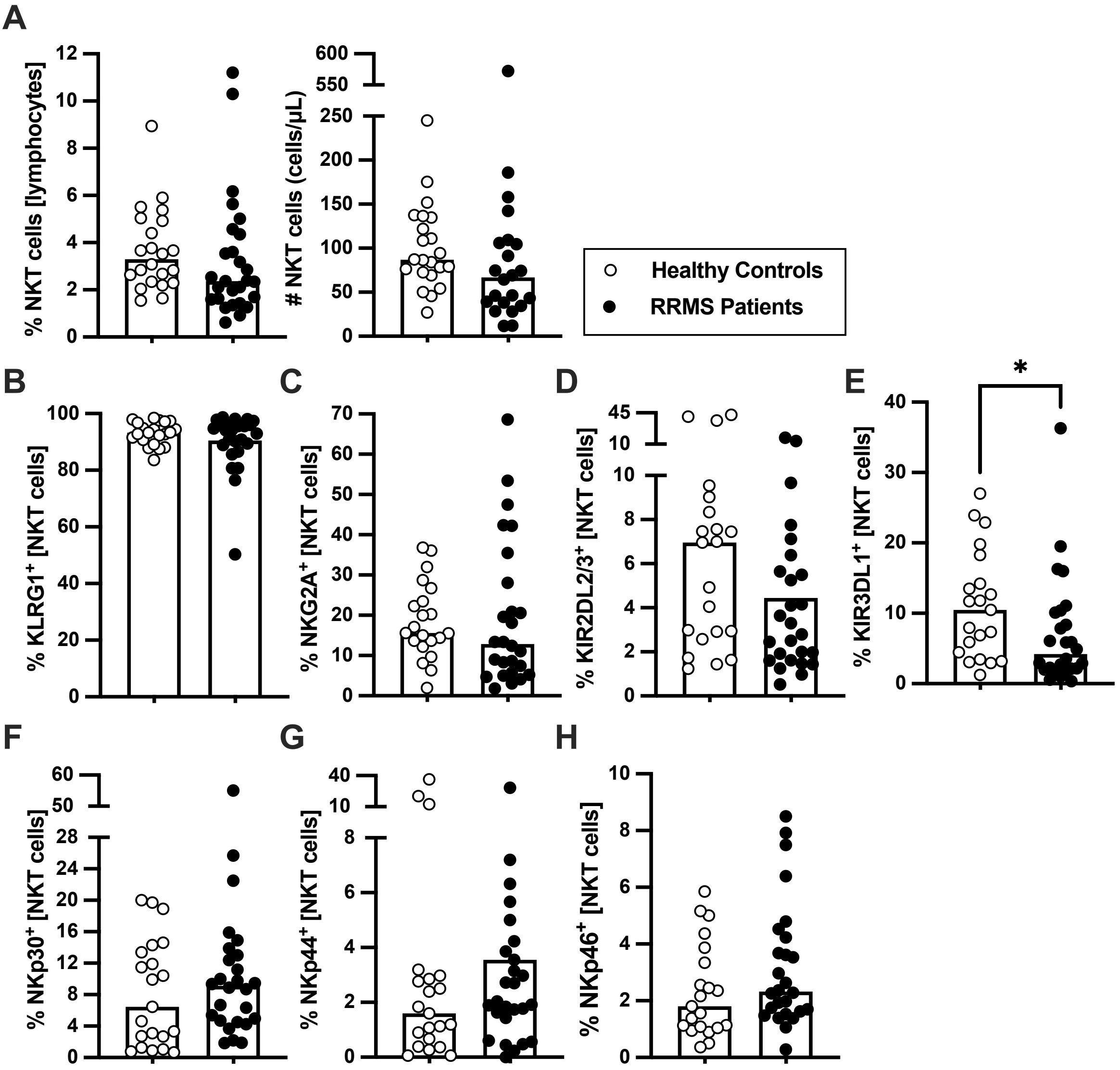

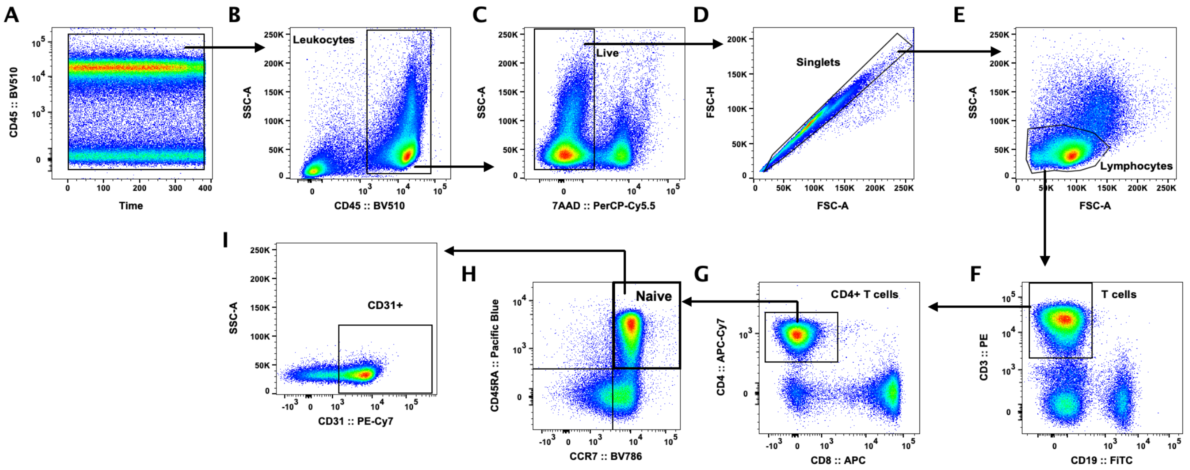

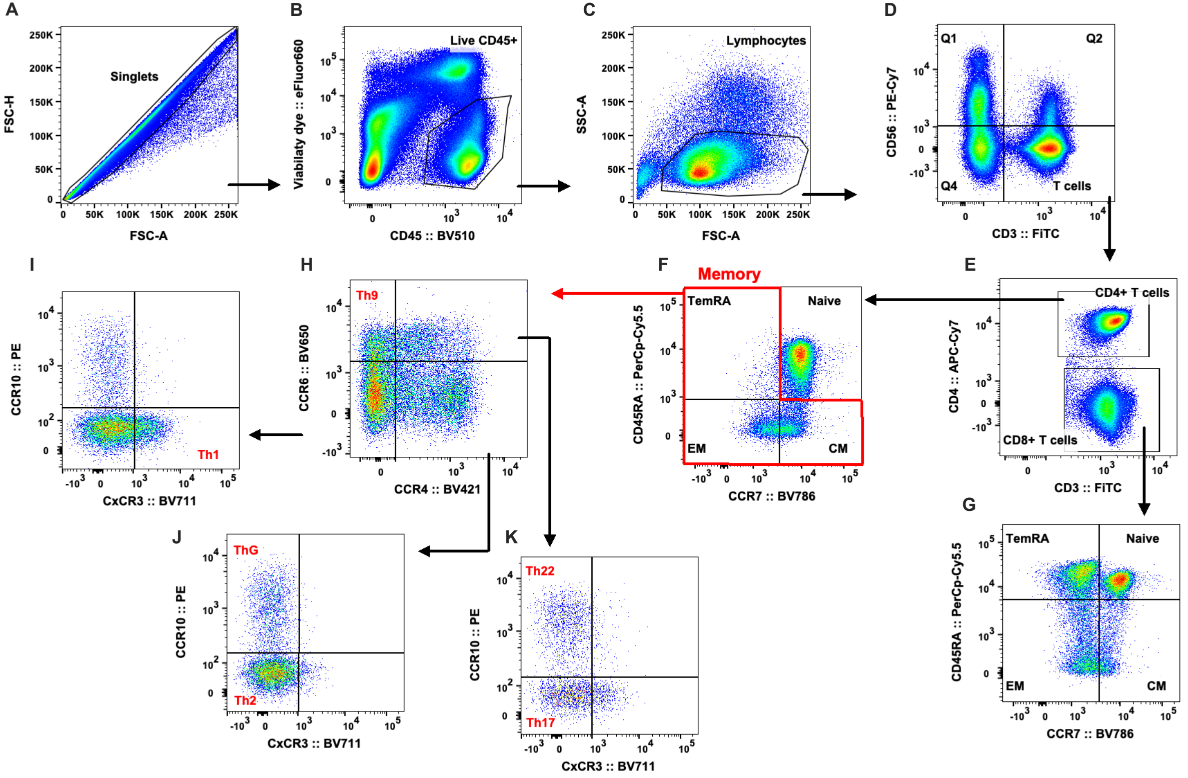

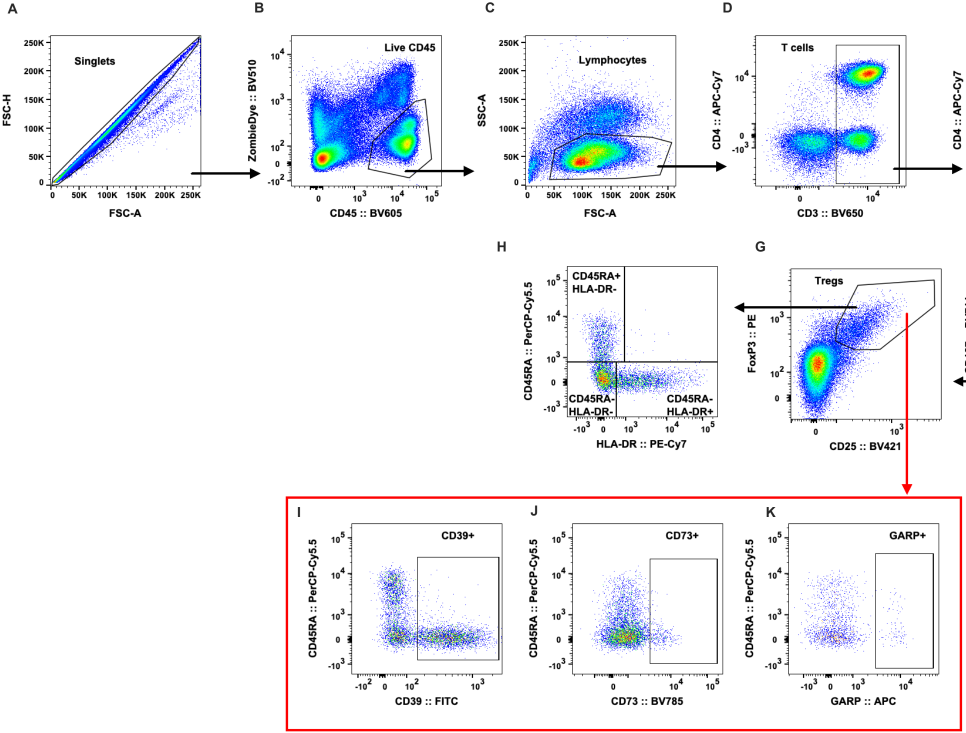

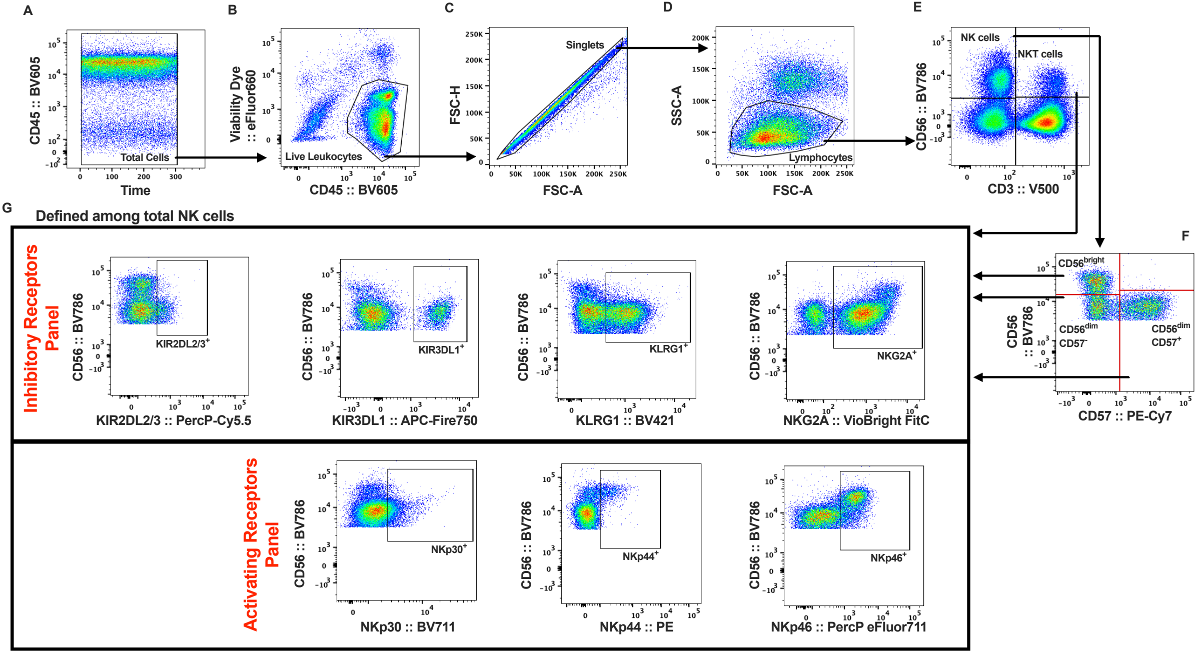
