## Supplemental Methods for "Low memory T cells blood counts and high naïve regulatory T cells percentage at relapsing remitting multiple sclerosis diagnosis"

### eMETHODS

#### Study Population

Patients included in this cross-sectional study were followed-up between 2018 and 2021 as outpatients at *Hospital de Braga* (Braga, Portugal), *Hospital Geral de Santo António* (*Centro Hospitalar Universitário do Porto*, Porto, Portugal) and *Hospital Álvaro Cunqueiro* (Vigo, Spain). Patients older than 18 years old, newly diagnosed with RRMS according to McDonald 2017 criteria and naïve for MS disease modifying drugs were recruited. Exclusion criteria: other immune disorders (other autoimmune or immunodeficiency diseases); history of radiotherapy and/or chemotherapy; on immunosuppressive drugs; treated with corticosteroids on the last 3 months or continuously for more than 6 months; splenectomy or thymectomy; and/or pregnancy. Data on age, sex, age of MS onset, months living with MS, months from last relapse and disease severity [multiple sclerosis severity score (MSSS)] were retrieved from medical records (**eTable 12**). Healthy controls, age and sex-matched, were recruited at *Centro Clínico Académico, Hospital de Braga*.

#### Sample Processing

Blood was collected into $K_3$EDTA tubes. Processing of blood samples from the different hospitals was centralized at the Life and Health Sciences Research Institute (ICVS; Braga, Portugal) and performed, in the exact same conditions for all samples, within 6 hours after collection.

T cells and $CD4^+$ T cells absolute numbers were determined in 10 μL of whole blood through: i) Muse Human CD4 T cell Kit (Millipore Corporation, CA, USA), according to manufacturer's protocol, and acquired on a MUSE cell analyzer (Millipore Corporation); or ii) FACS staining with titrated anti-human PE-conjugated anti-CD3 (clone OKT3) and APC-Cy7-conjugated anti-CD4 (clone RPA-T4; both from BioLegend, CA, USA) for 15 min, followed by a 15 min incubation with red blood cells lysis buffer (Ammonium-Chloride-Potassium); prior to acquisition on a BD LSRII cytometer; a known number of counting beads (Molecular Probes, Oregon, USA) were added to each tube.

Peripheral blood mononuclear cells (PBMCs) and plasma were obtained from whole blood by Histopaque-1077 (Sigma-Aldrich, Missouri, USA) gradient centrifugation and enumerated. Plasma was aliquoted and frozen at -80 ºC. One million PBMCs were centrifuged at 13000 rpm and frozen as dry pellet for quantification of T cell receptor excision circles (TRECs). The remaining PBMCs were frozen, in 5 million cells aliquots, in RPMI media supplemented with 20% FBS (Pan Biotech, Aidenbach, Germany) and 10% dimethyl sulfoxide (DMSO; Sigma-Aldrich, Missouri, USA) and kept at -80ºC within a MrFrosty for at least 24 h prior being transferred to liquid nitrogen.

**Anti-Human Cytomegalovirus IgG seroprevalence**

Semi-qualitative measurement of plasma IgG antibodies against the human cytomegalovirus (HCMV) was performed using the anti-HCMV IgG Elisa kit (Abcam, Cambridge, UK), according to manufacturer's instructions.

**FACS staining**

PBMCs aliquots were thawed in a water bath and washed using FACS buffer (PBS with 2% BSA and 0.01% sodium azide). One million PBMCs per stain were incubated with the following antibody panels (**eTable 13**): i) T cell Recent Thymic Emigrants (RTEs); ii) $CD4^+$ and $CD8^+$ T cells subsets homeostasis; iii) Tregs characterization; and iv) inhibitory and activating NK/NKT cells study. After incubation for 20 min, RT in the dark, cells were thoroughly washed with FACS buffer or PBS. The RTEs homeostasis panel was acquired on the same day after incubation with 7AAD for 10 min. Cells from the remaining panels were incubated with a fixable viability dye (30 min, RT, dark), and fixed using the Fixation/Permeabilization solution from the FOXP3 Staining Buffer Set (eBiosciences, CA, USA). Cells from the Tregs stain were further permeabilized using the permeabilization buffer from the same kit followed by intracellular staining of FOXP3 (30 min, RT, dark). For the $CD4^+$ and $CD8^+$ T cells subsets homeostasis and Tregs characterization panels, all samples were analyzed in a single batch. As the samples for the RTE and NK/NKT cells panel were analyzed in several batches, a reference control consisting of PBMCs from one individual was always stained, acquired, and analyzed together with study samples to control for experimental and analysis variability. The Coefficient of Variation (CV) of the reference control sample was extracted for each cell population; the mean CV of all populations was 7.4 %, with the lowest CV of 0.1% for the percentage of KLRG1$^+$ NK cells, and the highest CV of 18.9 % for the percentage of NKp30$^+$ NK cells. Moreover, prior to acquisitions, rainbow calibration beads (BioLegend, CA, USA) were acquired, and voltages adjusted to maintain cytometer settings standardization between different experimental batches.

Samples were acquired on a BD LSRII flow cytometer, using the FACS Diva Software v6.0 (Becton Dickinson, NJ, USA). Data were analyzed in a blinded way, using the FlowJo Software v10 (Becton Dickinson, NJ, USA), as described in **eFigures 4, 5, 6 and 7**. Tregs subsets, and inhibitory and activating receptors expression among NK and NKT cells were evaluated only when parent population had at least 500 events.

**T Cell Receptor Excision Circles quantification**

Single joint (sj) and Dbeta-Jbeta (DJβ) T cell receptor excision circles (TRECs) were quantified through nested polymerase chain reaction (PCR), using primers and standard curve's plasmids, as previously described.[1] Briefly, dry pellets of 1 million PBMCs were thawed (10 min, RT) resuspended in 0.1 mL of lysis solution [10 mM Tris-HCl pH 8.0, Tween-20 (0.05%), 0.05% nonidet P-40 (NP-40 from Calbiochem; Merck KGaA, Darmstadt, Germany), and 100 µg/mL Proteinase K (Grisp, Porto, Portugal)], and incubated on a thermocycler (Mastercycler epgradient S; Eppendorf, Hamburg, Germany) for 30 min at 56 ºC, followed by 10 min at 98 ºC. Samples were kept at 4 ºC until being tested. sjTRECs and DJβTRECs were quantified in distinct assays, each together with CD3; two plasmids were used for quantification, each containing the sjTRECs or DJβTRECs sequence and the housekeeping gene CD3 at a 1:1 ratio.[1] The first round of amplification was performed on cell lysates and plasmid through a multiplex PCR using specific 5'/3' "out" primers to sjTRECs/CD3 or DJβTRECs/CD3 (**eTable 14**) and 0.5 U Xpert Taq DNA Polymerase, 2.5 mM $MgCl_2$ and 0.4 mM dNTPs mix (all from Grisp): 15 min at 95 ºC followed by 19 amplification cycles of 30 sec at 95 ºC, 30 sec at 60 ºC and 3 min at 72 ºC. The sample products of the first amplification were 10 or 50-fold diluted for sjTRECs/CD3 or DJβTRECs/CD3, respectively; the standard curve was prepared through 8 sequential 10-fold dilutions of the plasmid 1st amplification product. Using specific 5'/3' "in" primers (**eTable 14**) and 1x Kapa SYBR Fast qPCR Master Mix (Roche, Basel, Switzerland), TRECs and CD3 were amplified in independent wells, each in duplicate in the same qPCR run using either a Bio-Rad CFX96 real-time system with a C1000 thermal cycler (Bio-Rad, CA, USA) or a Applied Biosystems 7500 Real-Time PCR Systems (Thermo Fisher Scientific, Massachusetts, USA): 15 min at 95 ºC for, followed by 40 cycles of 2 sec at 95 ºC, 15 sec at 62 or 60 ºC for sjTREC/CD3 or DJβTRECs/CD3, respectively, and 10 sec at 72 ºC. The fluorescence signal was measured in each well at the end of each cycle. Melting curves were analyzed and showed a single sharp peak at the temperature characteristic of the primers used. Data were analyzed in Bio-Rad CFX Manager v3.1 (Bio-Rad) or Design and Analysis Software v2.4.3 (Thermo Fisher Scientific) and sjTREC and DJβTRECs quantification represented as number of TRECs/$10^5$ cells. The sj/DJβTREC ratio was calculated as previously described.[1]

**Statistical analysis**

Statistical analysis was performed using IBM SPSS Statistics v26 (IBM Corporation, NY, USA) and GraphPad Prism v9 (GraphPad Software, CA, USA). Differences were considered statistically significant for *p*-values <0.050. The histogram and measures of asymmetry and kurtosis were evaluated, and the D'Agostino & Pearson test performed to assess the normality assumption for parametric tests; for data not following a normal distribution, non-parametric tests were used. For quantitative data, depending on the underlying distributions, comparisons between two independent groups were performed through independent *t*-test or Mann-Whitney *U*-test. For qualitative data, comparison between two groups was performed through the Chi-square test. As a measure of the magnitude of a difference, the effect size (practical significance) was calculated as follows[2]:

- *t*-tests, as the groups have unequal sample sizes, Cohen's *d* was calculated as:

$$d = \frac{M_1 - M_2}{\sqrt{\frac{(N_1 - 1)SD_1^2 + (N_2 - 1)SD_2^2}{(N_1 - N_2)}}}$$

where:

- $M_1$, $M_2$: Mean of group 1 and group 2, respectively.
- $N_1$, $N_2$: Sample size of group 1 and group 2, respectively.
- $SD_1$, $SD_2$: Standard deviation of group 1 and group 2, respectively.

Cohen's *d* was considered small if $d$ <0.300; medium if $d$ = [0.300,0.800]; or large if $d$ >0.800.

- Mann-Whitney *U*-test, *r* was calculated as:

$$r = \frac{Z}{\sqrt{N}}$$

where:

- Z: z-score.
- N: Total sample size.

r was considered small if *r* <0.300, medium if *r* = [0.300,0.500], or large if *r* >0.500. Sample size, statistical outputs and effect size are summarized in **eTable 1.** As sex, age and HCMV IgG seroprevalence contribute to variations on immune cells percentages and numbers[3–5], all comparisons were confirmed through multiple linear regression models to adjust for these variables. The dependent variable was the cell population in study (either frequency or number), and the independent variables age, sex (reference category: female), HCMV IgG seroprevalence (reference: IgG negative) and RRMS (reference: healthy). All differences between study groups were maintained in those models (**eTables 2-9**), unless otherwise stated in the results section. MS clinical parameters contribution to immune cell populations alterations were evaluated through sequential multiple linear regressions followed by simple linear regressions. First, to control for age, sex, and HCMV IgG seroprevalence multiple linear regressions having as dependent variable each of the cell populations were performed, and the unstandardized residuals retrieved.[6] Then, the predictive value of each MS clinical parameter was evaluated through simple linear regression having as dependent variable the abovementioned unstandardized residuals for each cell population, and as independent variables either the time from the last relapse or the MS severity score (**eTables 10, 11**). Corticoids' administration was included also as an independent dichotomic variable in the regression for time from last relapse to control for the possible residual consequences of corticosteroids administration for relapse treatment. Effect size $R^2$ for the linear regression was considered small, medium, and large if $R^2$ <0.090, $R^2$ = [0.090;0.250] and $R^2$ >0.250, respectively.


1. Dion, M., Sékaly, R. & Cheynier, R. Estimating thymic function through quantification of T-cell receptor excision circles. *Methods Mol. Biol.* **380**, 197–213 (2007).
2. Cohen, J. *Statistical power analysis for the behavioral sciences*. (Lawrence Erlbaum Associates, 1988).
3. Márquez, E. J. *et al.* Sexual-dimorphism in human immune system aging. *Nat. Commun.* **11**, (2020).
4. Giefing-Kröll, C., Berger, P., Lepperdinger, G. & Grubeck-Loebenstein, B. How sex and age affect immune responses, susceptibility to infections, and response to vaccination. *Aging Cell* **14**, 309–321 (2015).
5. Sheikh, M. H. *et al.* Immuno-metabolic impact of the multiple sclerosis patients' sera on endothelial cells of the blood-brain barrier. *J. Neuroinflammation* **17**, 1–19 (2020).
6. Marques, P. *et al.* The functional connectome of cognitive reserve. *Hum. Brain Mapp.* **37**, 3310–3322 (2016).
