## Supplemental Tables for "Low memory T cells blood counts and high naïve regulatory T cells percentage at relapsing remitting multiple sclerosis diagnosis"

**eTable 1.** Statistical outputs and sample size calculations.

| | Percentage of cells | | | | | Cells absolute numbers | | | | | Fig. |
|---|---|---|---|---|---|---|---|---|---|---|---|
| | Sample Size | | Test result | *p-value*[1] | Effect size[2] | Sample Size | | Test result | *p-value*[1] | Effect size[2] | |
| | HC | RRMS | | | | HC | RRMS | | | | |
| **CD31$^+$ naive CD4$^+$ T cells** | 20 | 19 | $U$ = 137,5 | 0,143 | 0,223 | 20 | 19 | $U$ = 163 | 0,628 | 0,065 | 1A |
| **sj/DJβTREC** | 16 | 16 | $t_{30}$ = 0,157 | 0,876 | 0,092 | | | | | | 1B |
| **sjTRECs** | 20 | 19 | $U$ = 139,0 | 0,155 | 0,225 | | | | | | 1C |
| **DJβTREC** | 16 | 16 | $U$ = 98,000 | 0,179 | 0,226 | | | | | | 1D |
| **CD4$^+$ T cells** | 33 | 30 | $t_{61}$ = 1,722 | 0,090 | 0,434 | 33 | 28 | $t_{59}$ = 1,413 | 0,163 | 0,361 | 2A |
| **Naive/Memory CD4$^+$ T cells** | 33 | 30 | $U$ = 413,0 | 0,263 | 0,141 | | | | | | 2C |
| **Naïve CD4$^+$ T cells** | 33 | 30 | $t_{61}$ = 0,969 | 0,336 | 0,244 | 33 | 28 | $U$ = 426,0 | 0,607 | 0,067 | 2D |
| **Memory CD4$^+$ T cells** | 33 | 30 | $t_{61}$ = 0,969 | 0,336 | 0,244 | 33 | 28 | $U$ = 356,0 | 0,127 | 0,196 | 2E |
| **CM CD4$^+$ T cells** | 33 | 30 | $t_{61}$ = 0,575 | 0,568 | 0,145 | 33 | 28 | $U$ = 392,5 | 0,319 | 0,130 | 2F |
| **EM CD4$^+$ T cells** | 33 | 30 | $U$ = 391 | 0,154 | 0,180 | 33 | 28 | $U$ = 351,500 | 0,111 | 0,204 | 2G |
| **TemRA CD4$^+$ T cells** | 33 | 30 | $U$ = 368,0 | 0,081 | 0,220 | 33 | 28 | $U$ = 337,5 | 0,072 | 0,228 | 2H |
| **CD8$^+$ T cells** | 33 | 30 | $U$ = 365,500 | 0,075 | 0,225 | 33 | 28 | $t_{59}$ = 2,647 | 0,010 | 0,682 | 3A |
| **Naive/Memory CD8$^+$ T cells** | 33 | 30 | $U$ = 366,0 | 0,076 | 0,225 | | | | | | 3C |
| **Naïve CD8$^+$ T cells** | 33 | 30 | $t_{61}$ = 2,151 | 0,035 | 0,543 | 33 | 28 | $t_{59}$ = 0,664 | 0,509 | 0,171 | 3D |
| **Memory CD8$^+$ T cells** | 33 | 30 | $t_{61}$ = 2,151 | 0,035 | 0,543 | 33 | 28 | $t_{59}$ = 3,325 | 0,002 | 0,859 | 3E |
| **CM CD8$^+$ T cells** | 33 | 30 | $t_{61}$ = 0,925 | 0,358 | 0,233 | 33 | 28 | $U$ = 364,5 | 0,160 | 0,176 | 3F |
| **EM CD8$^+$ T cells** | 33 | 30 | $U$ = 492 | 0,970 | 0,005 | 33 | 28 | $U$ = 311,000 | 0,028 | 0,280 | 3G |
| **TemRA CD8$^+$ T cells** | 33 | 30 | $U$ = 298,0 | 0,006 | 0,342 | 33 | 28 | $U$ = 230,5 | 0,001 | 0,428 | 3H |
| **Treg** | 33 | 30 | $U$ = 432,5 | 0,394 | 0,108 | 33 | 28 | $U$ = 434,0 | 0,690 | 0,056 | 4A |
| **Naive/Memory Tregs** | 33 | 30 | $U$ = 316,5 | 0,013 | 0,310 | | | | | | 4C |
| **CD45RA$^+$HLA-DR$^-$ Tregs** | 33 | 30 | $U$ = 317 | 0,014 | 0,309 | 33 | 28 | $t_{59}$ = 1,45 | 0,152 | 0,371 | 4D |
| **CD45RA$^-$HLA-DR$^-$ Tregs** | 33 | 30 | $t_{61}$ = 1,888 | 0,064 | 0,476 | 33 | 28 | $U$ = 367,0 | 0,171 | 0,178 | 4E |
| **CD45RA$^-$HLA-DR$^+$ Tregs** | 33 | 30 | $U$ = 381,5 | 0,120 | 0,197 | 33 | 28 | $U$ = 380,5 | 0,241 | 0,157 | 4F |
| **CD39$^+$ Tregs** | 33 | 30 | $t_{61}$ = 2,019 | 0,048 | 0,509 | 33 | 28 | $U$ = 354,0 | 0,120 | 0,200 | 4G |
| **CD73$^+$ Tregs** | 33 | 30 | $U$ = 464,000 | 0,674 | 0,054 | 33 | 28 | $U$ = 416,0 | 0,511 | 0,085 | 4H |
| **GARP$^+$ Tregs** | 33 | 30 | $U$ = 286,000 | 0,004 | 0,362 | 33 | 28 | $U$ = 289,0 | 0,012 | 0,321 | 4I |
| **NK cells** | 22 | 27 | $U$ = 295,5 | 0,980 | 0,004 | 22 | 22 | $U$ = 215,0 | 0,538 | 0,096 | 5A |
| **CD56$^{bright}$ NK cells** | 22 | 27 | $U$ = 247,0 | 0,320 | 0,144 | 22 | 22 | $t_{42}$ = 1,156 | 0,254 | 0,349 | 5B |
| **CD56$^{dim}$CD57$^-$ NK cells** | 22 | 27 | $t_{47}$ = 0,913 | 0,366 | 0,262 | 22 | 22 | $U$ = 237,0 | 0,917 | 0,018 | 5C |
| **CD56$^{dim}$CD57$^+$ NK cells** | 22 | 27 | $t_{47}$ = 1,329 | 0,190 | 0,382 | 22 | 22 | $U$ = 215,0 | 0,538 | 0,096 | 5D |
| **KLRG1$^+$CD56$^{bright}$ NK cells** | 17 | 22 | $U$ = 215,5 | 0,006 | 0,262 | | | | | | 5E |
| **KLRG1$^+$ CD56$^{dim}$CD57$^-$ NK cells** | 21 | 26 | $t_{45}$ = 0,040 | 0,969 | 0,012 | | | | | | |
| **KLRG1$^+$ CD56$^{dim}$CD57$^+$ NK cells** | 21 | 26 | $t_{45}$ = 1,072 | 0,289 | 0,315 | | | | | | |
| **NKG2A$^+$ CD56$^{bright}$ NK cells** | 17 | 22 | $U$ = 185,5 | 0,056 | 0,359 | | | | | | 5F |
| **NKG2A$^+$ CD56$^{dim}$CD57$^-$ NK cells** | 21 | 26 | $t_{45}$ = 1,832 | 0,074 | 0,538 | | | | | | |
| **NKG2A$^+$ CD56$^{dim}$CD57$^+$ NK cells** | 21 | 26 | $t_{45}$ = 1,994 | 0,052 | 0,585 | | | | | | |
| **KIR2DL2/3$^+$ CD56$^{bright}$ NK cells** | 17 | 22 | $t_{37}$ = 0,187 | 0,852 | 0,061 | | | | | | 5G |
| **KIR2DL2/3$^+$ CD56$^{dim}$CD57$^-$ NK cells** | 21 | 26 | $t_{45}$ = 0,427 | 0,672 | 0,125 | | | | | | |
| **KIR2DL2/3$^+$ CD56$^{dim}$CD57$^+$ NK cells** | 21 | 26 | $U$ = 280,5 | 0,795 | 0,048 | | | | | | |
| **KIR3DL1$^+$ CD56$^{bright}$ NK cells** | 17 | 22 | $U$ = 293,0 | 0,732 | 0,013 | | | | | | 5H |
| **KIR3DL1$^+$ CD56$^{dim}$CD57$^-$ NK cells** | 21 | 26 | $U$ = 248,0 | 0,320 | 0,144 | | | | | | |
| **KIR3DL1$^+$ CD56$^{dim}$CD57$^+$ NK cells** | 21 | 26 | $U$ = 238,0 | 0,235 | 0,173 | | | | | | |
| **NKp30$^+$ CD56$^{bright}$ NK cells** | 17 | 15 | $U$ = 243,0 | 0,143 | 0,192 | | | | | | 5I |
| **NKp30$^+$ CD56$^{dim}$CD57$^-$ NK cells** | 22 | 27 | $t_{47}$ = 0,135 | 0,893 | 0,038 | | | | | | |
| **NKp30$^+$ CD56$^{dim}$CD57$^+$ NK cells** | 22 | 26 | $U$ = 162,0 | 0,009 | 0,392 | | | | | | |
| **NKp44$^+$ CD56$^{bright}$ NK cells** | 17 | 15 | $U$ = 193,5 | 0,395 | 0,368 | | | | | | 5J |
| **NKp44$^+$ CD56$^{dim}$CD57$^-$ NK cells** | 22 | 27 | $U$ = 234,0 | 0,209 | 0,181 | | | | | | |
| **NKp44$^+$ CD56$^{dim}$CD57$^+$ NK cells** | 22 | 26 | $U$ = 278,0 | 0,626 | 0,055 | | | | | | |
| **NKp46$^+$ CD56$^{bright}$ NK cells** | 17 | 15 | $t_{30}$ = 0,294 | 0,771 | 0,109 | | | | | | 5K |
| **NKp46$^+$ CD56$^{dim}$CD57$^-$ NK cells** | 22 | 27 | $U$ = 247,0 | 0,320 | 0,144 | | | | | | |
| **NKp46$^+$ CD56$^{dim}$CD57$^+$ NK cells** | 22 | 26 | $t_{46}$ = 1,035 | 0,306 | 0,301 | | | | | | |
| **Th1 CD4$^+$ T cells** | 33 | 30 | $U$ = 439,500 | 0,450 | 0,095 | 33 | 28 | $U$ = 439,000 | 0,747 | 0,043 | e1A |
| **Th2 CD4$^+$ T cells** | 33 | 30 | $t_{61}$ = 0,9254 | 0,358 | 0,233 | 33 | 28 | $U$ = 349,0 | 0,104 | 0,209 | e1B |
| **Th9 CD4$^+$ T cells** | 33 | 30 | $U$ = 463 | 0,664 | 0,055 | 33 | 28 | $U$ = 443,0 | 0,790 | 0,035 | e1C |
| **Th17 CD4$^+$ T cells** | 33 | 30 | $U$ = 479,5 | 0,835 | 0,029 | 33 | 28 | $U$ = 445,5 | 0,816 | 0,030 | e1D |
| **Th22 CD4$^+$ T cells** | 33 | 30 | $U$ = 475,5 | 0,792 | 0,040 | 33 | 28 | $U$ = 454 | 0,911 | 0,015 | e1E |
| **ThG CD4$^+$ T cells** | 33 | 30 | $U$ = 474,5 | 0,782 | 0,035 | 33 | 28 | $U$ = 401,000 | 0,384 | 0,113 | e1F |

### eTable 1 (cont)

| | Percentage of cells | | | | | Cells absolute numbers | | | | | Fig. |
|---|---|---|---|---|---|---|---|---|---|---|---|
| | Sample Size | | Test result | *p-value*[1] | Effect size[2] | Sample Size | | Test result | *p-value*[1] | Effect size[2] | |
| | HC | RRMS | | | | HC | RRMS | | | | |
| **CCR4⁺ Memory CD8⁺ T cells** | 33 | 30 | *U* = 479,500 | 0,835 | 0,027 | 33 | 28 | *U* = 364 | 0,159 | 0,182 | e2A |
| **CCR6⁺ Memory CD8⁺ T cells** | 33 | 30 | *U* = 460,000 | 0,635 | 0,061 | 33 | 28 | *U* = 321 | 0,041 | 0,261 | |
| **CCR10⁺ Memory CD8⁺ T cells** | 33 | 30 | *U* = 426,000 | 0,347 | 0,120 | 33 | 28 | *U* = 410 | 0,459 | 0,015 | |
| **CxCR3⁺ Memory CD8⁺ T cells** | 33 | 30 | *U* = 398,500 | 0,187 | 0,167 | 33 | 28 | *U* = 454 | 0,914 | 0,096 | |
| **CCR4⁺ CM CD8⁺ T cells** | 33 | 30 | *U* = 404,500 | 0,216 | 0,157 | 33 | 28 | *U* = 372,500 | 0,198 | 0,165 | e2B |
| **CCR6⁺ CM CD8⁺ T cells** | 33 | 30 | $t_{61}$ = 0,937 | 0,352 | 0,237 | 33 | 28 | *U* = 394,0 | 0,329 | 0,124 | |
| **CCR10⁺ CM CD8⁺ T cells** | 33 | 30 | *U* = 442,000 | 0,473 | 0,092 | 33 | 28 | *U* = 401,5 | 0,386 | 0,022 | |
| **CxCR3⁺ CM CD8⁺ T cells** | 33 | 30 | $t_{61}$ = 1,248 | 0,217 | 0,315 | 33 | 28 | *U* = 450 | 0,869 | 0,111 | |
| **CCR4⁺ EM CD8⁺ T cells** | 33 | 30 | *U* = 460,000 | 0,635 | 0,061 | 33 | 28 | *U* = 377 | 0,223 | 0,098 | e2C |
| **CCR6⁺ EM CD8⁺ T cells** | 33 | 30 | *U* = 337,000 | 0,030 | 0,274 | 33 | 28 | *U* = 409 | 0,450 | 0,157 | |
| **CCR10⁺ EM CD8⁺ T cells** | 33 | 30 | *U* = 447,500 | 0,518 | 0,082 | 33 | 28 | *U* = 425 | 0,600 | 0,024 | |
| **CxCR3⁺ EM CD8⁺ T cells** | 33 | 30 | *U* = 444,500 | 0,492 | 0,087 | 33 | 28 | *U* = 449 | 0,855 | 0,069 | |
| **CCR4⁺ TemRA CD8⁺ T cells** | 33 | 30 | *U* = 489,000 | 0,937 | 0,010 | 33 | 28 | *U* = 337,5 | 0,072 | 0,230 | e2D |
| **CCR6⁺ TemRA CD8⁺ T cells** | 33 | 30 | $t_{61}$ = 2,748 | 0,008 | 0,693 | 33 | 28 | *U* = 199 | 0,000 | 0,487 | |
| **CCR10⁺ TemRA CD8⁺ T cells** | 33 | 30 | *U* = 384,000 | 0,128 | 0,193 | 33 | 28 | *U* = 352,5 | 0,114 | 0,084 | |
| **CxCR3⁺ TemRA CD8⁺ T cells** | 33 | 30 | *U* = 374,500 | 0,098 | 0,209 | 33 | 28 | *U* = 418 | 0,529 | 0,202 | |
| **NKT cells** | 22 | 27 | *U* = 213,5 | 0,095 | 0,240 | 22 | 22 | *U* = 165,0 | 0,072 | 0,272 | e3A |
| **KLRG1⁺ NKT cells** | 21 | 26 | *U* = 270,0 | 0,845 | 0,079 | | | | | | e3B |
| **NKG2A⁺ NKT cells** | 21 | 26 | *U* = 257,0 | 0,353 | 0,117 | | | | | | e3C |
| **KIR2DL2/3⁺ NKT cells** | 21 | 26 | *U* = 218,5 | 0,071 | 0,230 | | | | | | e3D |
| **KIR3DL1⁺ NKT cells** | 21 | 26 | *U* = 195,0 | 0,015 | 0,299 | | | | | | e3E |
| **NKp30⁺ NKT cells** | 21 | 26 | *U* = 263,0 | 0,344 | 0,100 | | | | | | e3F |
| **NKp44⁺ NKT cells** | 21 | 26 | *U* = 263,0 | 0,320 | 0,100 | | | | | | e3G |
| **NKp46⁺ NKT cells** | 21 | 26 | *U* = 223,5 | 0,128 | 0,215 | | | | | | e3H |

[1] Red cells refere to significant differences (*p*-value <0.050) and grey cells to tendencies (0.050< *p*-value <0.010)

[2] Cohen's *d* was calculated for *t*-tests, and *r* for Mann-Witney *U*-tests. Green cells refere to large effect sizes (*d*>0.800; *r*>0.500) and yellow cells to medium effect sizes (0.300≤*d*≤0.800; 0.300≤*r*≤0.500)

**CM**, Central Memory; **EM**, Effector Memory; **NK**, Natural Killer; **NKT**, Natural Killer T cells; **TemRA**, Terminally differentiated CD45RA-expressing memory cells; **Th**, T helper cells; **TREC**, T cell receptor excision circles; **Treg**, Regulatory T cells.

**eTable 2.** Multiple linear regression models to evaluate the impact of sex (female *vs.* male), age, HCMV IgG seroprevalence and disease status (healthy *vs.* RRMS) on thymic function surrogates.

| Dependent Variable | Independent Variable | Percentage of cells: Model | *p-value* | B | SE | *β* | *t* | p-value | Absolute number of cells: Model | *p-value* | B | SE | *β* | *t* | p-value | Fig |
|---|---|---|---|---|---|---|---|---|---|---|---|---|---|---|---|---|
| **$CD31^{+}$ Naive $CD4^{+}$ T cells** | | F(4,34)= 12,481 | **0,000** | | | | | | F(4,34)= 2,774 | **0,043** | | | | | | 1A |
| | Sex[1] | | | **-30,614** | **7,245** | **-0,580** | **-4,225** | **0,000** | | | -165,929 | 143,998 | -0,219 | -1,152 | 0,257 | |
| | Age | | | -0,357 | 0,284 | -0,177 | -1,257 | 0,217 | | | -9,738 | 5,762 | -0,327 | -1,690 | 0,100 | |
| | HCMV IgG[2] | | | **9,278** | **4,262** | **0,253** | **2,177** | **0,037** | | | 84,814 | 86,062 | 0,155 | 0,986 | 0,332 | |
| | RRMS[3] | | | -5,485 | 3,670 | -0,171 | -1,495 | 0,144 | | | 5,567 | 73,426 | 0,012 | 0,076 | 0,940 | |
| **sj/DJβTREC** | | F(4,28)= 0,926 | 0,463 | | | | | | | | | | | | | 1B |
| | Sex[1] | | | -6,978 | 9,022 | -0,176 | -0,773 | 0,446 | | | | | | | | |
| | Age | | | -0,265 | 0,370 | -0,168 | -0,716 | 0,480 | | | | | | | | |
| | HCMV IgG[2] | | | -3,735 | 5,825 | -0,123 | -0,641 | 0,527 | | | | | | | | |
| | RRMS[3] | | | -1,109 | 5,024 | -0,043 | -0,221 | 0,827 | | | | | | | | |
| **sjTREC[4]** | | F(4,34)= 7,866 | **0,000** | | | | | | | | | | | | | 1C |
| | Sex[1] | | | -141,369 | 139,469 | -0,126 | -1,014 | 0,316 | | | | | | | | |
| | Age | | | **-30,510** | **6,124** | **-0,603** | **-4,982** | **0,000** | | | | | | | | |
| | HCMV IgG[2] | | | -40,483 | 127,629 | -0,039 | -0,317 | 0,753 | | | | | | | | |
| | RRMS[3] | | | -69,328 | 114,685 | -0,076 | -0,605 | 0,549 | | | | | | | | |
| **DJβTREC[4]** | | F(4,28)= 2,627 | 0,056 | | | | | | | | | | | | | 1D |
| | Sex[1] | | | 6,157 | 22,742 | 0,056 | 0,271 | 0,789 | | | | | | | | |
| | Age | | | **-2,189** | **0,932** | **-0,501** | **-2,349** | **0,026** | | | | | | | | |
| | HCMV IgG[2] | | | -4,600 | 14,682 | -0,055 | -0,313 | 0,756 | | | | | | | | |
| | RRMS[3] | | | -10,597 | 12,663 | -0,147 | -0,837 | 0,410 | | | | | | | | |

[1] Reference is "Female"; [2] Reference is "IgG negative"; [3] Reference is "Healthy"; [4] Evaluated only when at leats 10 copies were detected

**HCMV,** Human Cytomegalovirus; **TREC**, T cell receptor excision circles.

**eTable 3.** Multiple linear regression models to evaluate the impact of sex (female *vs.* male), age, HCMV IgG seroprevalence and disease status (healthy *vs.* RRMS) on CD4$^{+}$ T cell subsets.

| Dependent Variable | Independent Variable | Percentage of cells: Model | *p-value* | B | SE | β | t | *p*-value | Absolute number of cells: Model | *p-value* | B | SE | β | t | *p*-value | Fig |
|---|---|---|---|---|---|---|---|---|---|---|---|---|---|---|---|---|
| **CD4$^{+}$ T cells** | | F(4,58)= 2,804 | **0,034** | | | | | | F(4,56)= 1,799 | 0,142 | | | | | | 2A |
| | Sex[1] | | | **-5,963** | **2,378** | **-0,313** | **-2,507** | **0,015** | | | **-230,602** | **105,586** | **-0,285** | **-2,184** | **0,033** | |
| | Age | | | 0,262 | 0,135 | 0,247 | 1,937 | 0,058 | | | 6,088 | 5,930 | 0,136 | 1,027 | 0,309 | |
| | HCMV IgG[2] | | | -1,624 | 2,673 | -0,077 | -0,607 | 0,546 | | | 24,684 | 120,488 | 0,027 | 0,205 | 0,838 | |
| | RRMS[3] | | | 3,619 | 2,279 | 0,197 | 1,588 | 0,118 | | | -140,812 | 100,203 | -0,181 | -1,405 | 0,165 | |
| **Naive/memory CD4$^{+}$ T cells** | | F(4,58)= 3,603 | **0,011** | | | | | | | | | | | | | 2C |
| | Sex[1] | | | -0,269 | 0,279 | -0,118 | -0,962 | 0,340 | | | | | | | | |
| | Age | | | **-0,045** | **0,016** | **-0,356** | **-2,854** | **0,006** | | | | | | | | |
| | HCMV IgG[2] | | | -0,338 | 0,314 | -0,134 | -1,076 | 0,286 | | | | | | | | |
| | RRMS[3] | | | -0,005 | 0,268 | -0,002 | -0,019 | 0,985 | | | | | | | | |
| **Naive CD4$^{+}$ T cells** | | F(4,58)= 6,099 | **0,000** | | | | | | F(4,56)= 3,280 | **0,017** | | | | | | 2D |
| | Sex[1] | | | **-7,157** | **3,474** | **-0,236** | **-2,060** | **0,044** | | | **-229,267** | **84,188** | **-0,340** | **-2,723** | **0,009** | |
| | Age | | | **-0,696** | **0,198** | **-0,412** | **-3,520** | **0,001** | | | -6,628 | 4,729 | -0,178 | -1,402 | 0,167 | |
| | HCMV IgG[2] | | | -0,751 | 3,905 | -0,022 | -0,192 | 0,848 | | | 35,183 | 96,070 | 0,046 | 0,366 | 0,716 | |
| | RRMS[3] | | | 3,461 | 3,329 | 0,118 | 1,040 | 0,303 | | | -60,447 | 79,896 | -0,093 | -0,757 | 0,452 | |
| **Memory CD4$^{+}$ T cells** | | F(4,58)= 6,099 | **0,000** | | | | | | F(4,56)= 4,201 | **0,005** | | | | | | 2E |
| | Sex[1] | | | **7,157** | **3,474** | **0,236** | **2,060** | **0,044** | | | -1,335 | 60,931 | -0,003 | -0,022 | 0,983 | |
| | Age | | | **0,696** | **0,198** | **0,412** | **3,520** | **0,001** | | | **12,716** | **3,422** | **0,460** | **3,716** | **0,000** | |
| | HCMV IgG[2] | | | 0,751 | 3,905 | 0,022 | 0,192 | 0,848 | | | -10,499 | 69,530 | -0,018 | -0,151 | 0,881 | |
| | RRMS[3] | | | -3,461 | 3,329 | -0,118 | -1,040 | 0,303 | | | -80,365 | 57,824 | -0,167 | -1,390 | 0,170 | |
| **CM CD4$^{+}$ T cells** | | F(4,58)= 1,994 | 0,107 | | | | | | F(4,56)= 1,230 | 0,309 | | | | | | 2F |
| | Sex[1] | | | 2,270 | 1,898 | 0,153 | 1,196 | 0,236 | | | -26,189 | 39,685 | -0,088 | -0,660 | 0,512 | |
| | Age | | | **0,220** | **0,108** | **0,266** | **2,033** | **0,047** | | | **4,776** | **2,229** | **0,290** | **2,143** | **0,036** | |
| | HCMV IgG[2] | | | -1,729 | 2,133 | -0,105 | -0,811 | 0,421 | | | -42,222 | 45,286 | -0,124 | -0,932 | 0,355 | |
| | RRMS[3] | | | 0,670 | 1,818 | 0,047 | 0,369 | 0,714 | | | -4,755 | 37,661 | -0,017 | -0,126 | 0,900 | |
| **EM CD4$^{+}$ T cells** | | F(4,58)= 6,463 | **0,000** | | | | | | F(4,56)= 6,160 | **0,000** | | | | | | 2G |
| | Sex[1] | | | 4,352 | 2,190 | 0,226 | 1,987 | 0,052 | | | 23,745 | 32,038 | 0,086 | 0,741 | 0,462 | |
| | Age | | | **0,429** | **0,125** | **0,400** | **3,447** | **0,001** | | | **6,956** | **1,800** | **0,454** | **3,866** | **0,000** | |
| | HCMV IgG[2] | | | 0,750 | 2,461 | 0,035 | 0,305 | 0,762 | | | 10,761 | 36,560 | 0,034 | 0,294 | 0,770 | |
| | RRMS[3] | | | -3,712 | 2,098 | -0,200 | -1,769 | 0,082 | | | **-68,938** | **30,405** | **-0,258** | **-2,267** | **0,027** | |
| **TemRA CD4$^{+}$ T cells** | | F(4,58)= 3,775 | **0,009** | | | | | | F(4,56)= 4,234 | **0,005** | | | | | | 2H |
| | Sex[1] | | | 0,520 | 0,580 | 0,109 | 0,896 | 0,374 | | | 0,879 | 7,441 | 0,014 | 0,118 | 0,906 | |
| | Age | | | 0,047 | 0,033 | 0,178 | 1,437 | 0,156 | | | **0,989** | **0,418** | **0,292** | **2,366** | **0,021** | |
| | HCMV IgG[2] | | | **1,723** | **0,652** | **0,327** | **2,643** | **0,011** | | | **20,875** | **8,491** | **0,299** | **2,459** | **0,017** | |
| | RRMS[3] | | | -0,430 | 0,556 | -0,094 | -0,774 | 0,442 | | | -6,844 | 7,061 | -0,116 | -0,969 | 0,337 | |

[1] Reference is "Female"; [2] Reference is "IgG negative"; [3] Reference is "Healthy"

**HCMV,** Human Cytomegalovirus; **CM**, Central Memory; **EM**, Efector Memory; **TemRA**, Terminally differentiated CD45RA-expressing memory cells;

**eTable 4.** Multiple linear regression models to evaluate the impact of sex (female *vs.* male), age, HCMV IgG seroprevalence and disease status (healthy *vs.* RRMS) on T helper (Th) subsets.

| Dependent Variable | Independent Variable | Percentage of cells | | | | | | | Absolute number of cells | | | | | | | Fig |
|---|---|---|---|---|---|---|---|---|---|---|---|---|---|---|---|---|
| | | Model | *p*-value | B | SE | *β* | *t* | *p*-value | Model | *p*-value | B | SE | *β* | *t* | *p*-value | |
| **Th1** | | F(4,58)= 0,900 | 0,470 | | | | | | F(4,56)= 1,455 | 0,228 | | | | | | e1A |
| | Sex[1] | | | -1,518 | 0,874 | -0,230 | -1,738 | 0,088 | | | **-26,097** | **12,216** | **-0,282** | **-2,136** | **0,037** | |
| | Age | | | 0,049 | 0,050 | 0,134 | 0,989 | 0,327 | | | 0,981 | 0,686 | 0,192 | 1,429 | 0,159 | |
| | HCMV IgG[2] | | | 0,295 | 0,982 | 0,040 | 0,301 | 0,765 | | | 3,020 | 13,940 | 0,029 | 0,217 | 0,829 | |
| | RRMS[3] | | | 0,042 | 0,837 | 0,007 | 0,051 | 0,960 | | | -5,716 | 11,593 | -0,064 | -0,493 | 0,624 | |
| **Th2** | | F(4,58)= 6,790 | **0,000** | | | | | | F(4,56)= 4,028 | **0,006** | | | | | | e1B |
| | Sex[1] | | | **3,269** | **0,944** | **0,390** | **3,463** | **0,001** | | | 18,106 | 14,084 | 0,157 | 1,286 | 0,204 | |
| | Age | | | **0,142** | **0,054** | **0,303** | **2,637** | **0,011** | | | **2,387** | **0,791** | **0,375** | **3,018** | **0,004** | |
| | HCMV IgG[2] | | | -0,866 | 1,061 | -0,093 | -0,816 | 0,418 | | | -12,734 | 16,071 | -0,097 | -0,792 | 0,431 | |
| | RRMS[3] | | | -1,147 | 0,904 | -0,142 | -1,268 | 0,210 | | | -22,374 | 13,366 | -0,202 | -1,674 | 0,100 | |
| **Th9** | | F(4,58)= 0,902 | 0,469 | | | | | | F(4,56)= 2,041 | 0,101 | | | | | | e1C |
| | Sex[1] | | | -0,836 | 0,507 | -0,218 | -1,648 | 0,105 | | | **-16,815** | **6,796** | **-0,321** | **-2,474** | **0,016** | |
| | Age | | | 0,038 | 0,029 | 0,179 | 1,323 | 0,191 | | | 0,723 | 0,382 | 0,249 | 1,893 | 0,064 | |
| | HCMV IgG[2] | | | -0,265 | 0,570 | -0,063 | -0,465 | 0,643 | | | -5,880 | 7,755 | -0,098 | -0,758 | 0,452 | |
| | RRMS[3] | | | 0,039 | 0,486 | 0,011 | 0,080 | 0,937 | | | -4,544 | 6,449 | -0,090 | -0,705 | 0,484 | |
| **Th17** | | F(4,58)= 0,757 | 0,557 | | | | | | F(4,56)= 1,904 | 0,122 | | | | | | e1D |
| | Sex[1] | | | -0,376 | 0,278 | -0,180 | -1,353 | 0,181 | | | **-7,753** | **3,583** | **-0,282** | **-2,164** | **0,035** | |
| | Age | | | 0,013 | 0,016 | 0,114 | 0,841 | 0,404 | | | 0,255 | 0,201 | 0,168 | 1,267 | 0,210 | |
| | HCMV IgG[2] | | | -0,358 | 0,312 | -0,155 | -1,145 | 0,257 | | | -6,853 | 4,089 | -0,218 | -1,676 | 0,099 | |
| | RRMS[3] | | | -0,049 | 0,266 | -0,025 | -0,186 | 0,853 | | | -3,012 | 3,401 | -0,114 | -0,886 | 0,380 | |
| **Th22** | | F(4,58)= 1,789 | 0,143 | | | | | | F(4,56)= 2,292 | 0,071 | | | | | | e1E |
| | Sex[1] | | | **-0,509** | **0,214** | **-0,307** | **-2,381** | **0,021** | | | **-6,852** | **2,669** | **-0,330** | **-2,568** | **0,013** | |
| | Age | | | 0,002 | 0,012 | 0,020 | 0,155 | 0,877 | | | 0,044 | 0,150 | 0,038 | 0,294 | 0,770 | |
| | HCMV IgG[2] | | | -0,266 | 0,240 | -0,145 | -1,106 | 0,273 | | | -4,173 | 3,046 | -0,176 | -1,370 | 0,176 | |
| | RRMS[3] | | | -0,110 | 0,205 | -0,069 | -0,538 | 0,593 | | | -2,675 | 2,533 | -0,134 | -1,056 | 0,295 | |
| **ThG** | | F(4,58)= 1,437 | 0,233 | | | | | | F(4,56)= 2,808 | **0,034** | | | | | | e1F |
| | Sex[1] | | | -0,552 | 0,677 | -0,106 | -0,816 | 0,418 | | | -20,405 | 10,293 | -0,251 | -1,983 | 0,052 | |
| | Age | | | 0,064 | 0,038 | 0,221 | 1,659 | 0,103 | | | **1,221** | **0,578** | **0,272** | **2,113** | **0,039** | |
| | HCMV IgG[2] | | | -1,448 | 0,760 | -0,252 | -1,904 | 0,062 | | | **-26,612** | **11,745** | **-0,287** | **-2,266** | **0,027** | |
| | RRMS[3] | | | -0,684 | 0,648 | -0,137 | -1,055 | 0,296 | | | -14,462 | 9,768 | -0,185 | -1,481 | 0,144 | |

[1] Reference is "Female"; [2] Reference is "IgG negative"; [3] Reference is "Healthy"

**HCMV,** Human Cytomegalovirus; **Th**, T helper cells;

**eTable 5.** Multiple linear regression models to evaluate the impact of sex (female *vs.* male), age, HCMV IgG seroprevalence and disease status (healthy *vs.* RRMS) on CD8$^+$ T cell subsets.

| Dependent Variable | Independent Variable | Percentage of cells | | | | | | | Absolute number of cells | | | | | | | Fig. |
|---|---|---|---|---|---|---|---|---|---|---|---|---|---|---|---|---|
| | | Model | *p*-value | B | SE | *β* | *t* | *p*-value | Model | *p*-value | B | SE | *β* | *t* | *p*-value | |
| **CD8$^+$ T cells** | | F(4,58)= 2,816 | **0,033** | | | | | | F(4,56)= 2,173 | 0,084 | | | | | | 3A |
| | Sex[1] | | | **5,981** | **2,376** | **0,315** | **2,517** | **0,015** | | | 81,630 | 92,349 | 0,114 | 0,884 | 0,381 | |
| | Age | | | -0,263 | 0,135 | -0,248 | -1,944 | 0,057 | | | -6,106 | 5,187 | -0,154 | -1,177 | 0,244 | |
| | HCMV IgG[2] | | | 1,647 | 2,670 | 0,078 | 0,617 | 0,540 | | | 55,773 | 105,382 | 0,068 | 0,529 | 0,599 | |
| | RRMS[3] | | | -3,600 | 2,276 | -0,196 | -1,581 | 0,119 | | | **-211,645** | **87,641** | **-0,307** | **-2,415** | **0,019** | |
| **Naive/memory CD8$^+$ T cells** | | F(4,58)= 3,221 | **0,019** | | | | | | | | | | | | | 3C |
| | Sex[1] | | | -0,113 | 0,160 | -0,087 | -0,708 | 0,482 | | | | | | | | |
| | Age | | | **-0,021** | **0,009** | **-0,289** | **-2,293** | **0,025** | | | | | | | | |
| | HCMV IgG[2] | | | -0,065 | 0,180 | -0,046 | -0,363 | 0,718 | | | | | | | | |
| | RRMS[3] | | | **0,317** | **0,154** | **0,254** | **2,066** | **0,043** | | | | | | | | |
| **Naive CD8$^+$ T cells** | | F(4,58)= 4,059 | **0,006** | | | | | | F(4,56)= 1,276 | 0,290 | | | | | | 3D |
| | Sex[1] | | | -5,516 | 3,767 | -0,177 | -1,464 | 0,148 | | | -25,259 | 51,203 | -0,066 | -0,493 | 0,624 | |
| | Age | | | **-0,517** | **0,214** | **-0,297** | **-2,411** | **0,019** | | | -5,379 | 2,876 | -0,253 | -1,870 | 0,067 | |
| | HCMV IgG[2] | | | -0,225 | 4,233 | -0,007 | -0,053 | 0,958 | | | 35,440 | 58,429 | 0,081 | 0,607 | 0,547 | |
| | RRMS[3] | | | **7,959** | **3,609** | **0,265** | **2,205** | **0,031** | | | -23,734 | 48,592 | -0,064 | -0,488 | 0,627 | |
| **Memory CD8$^+$ T cells** | | F(4,58)= 4,059 | **0,006** | | | | | | F(4,56)= 3,539 | **0,012** | | | | | | 3E |
| | Sex[1] | | | 5,517 | 3,767 | 0,177 | 1,465 | 0,148 | | | 106,900 | 62,318 | 0,213 | 1,715 | 0,092 | |
| | Age | | | **0,517** | **0,214** | **0,297** | **2,411** | **0,019** | | | -0,727 | 3,500 | -0,026 | -0,208 | 0,836 | |
| | HCMV IgG[2] | | | 0,226 | 4,233 | 0,007 | 0,053 | 0,958 | | | 20,336 | 71,113 | 0,035 | 0,286 | 0,776 | |
| | RRMS[3] | | | **-7,960** | **3,609** | **-0,265** | **-2,206** | **0,031** | | | **-187,917** | **59,141** | **-0,388** | **-3,177** | **0,002** | |
| **CM CD8$^+$ T cells** | | F(4,58)= 2,661 | **0,042** | | | | | | F(4,56)= 1,133 | 0,351 | | | | | | 3F |
| | Sex[1] | | | 0,190 | 0,903 | 0,026 | 0,211 | 0,834 | | | 7,490 | 9,225 | 0,108 | 0,812 | 0,420 | |
| | Age | | | **0,143** | **0,051** | **0,357** | **2,788** | **0,007** | | | 0,705 | 0,518 | 0,185 | 1,360 | 0,179 | |
| | HCMV IgG[2] | | | -1,530 | 1,015 | -0,192 | -1,508 | 0,137 | | | -0,259 | 10,527 | -0,003 | -0,025 | 0,980 | |
| | RRMS[3] | | | 0,486 | 0,865 | 0,070 | 0,562 | 0,576 | | | -9,376 | 8,755 | -0,141 | -1,071 | 0,289 | |
| **EM CD8$^+$ T cells** | | F(4,58)= 1,809 | 0,139 | | | | | | F(4,56)= 2,555 | **0,049** | | | | | | 3G |
| | Sex[1] | | | 6,045 | 3,164 | 0,246 | 1,911 | 0,061 | | | **93,825** | **41,703** | **0,287** | **2,250** | **0,028** | |
| | Age | | | 0,161 | 0,180 | 0,117 | 0,892 | 0,376 | | | -1,781 | 2,342 | -0,099 | -0,760 | 0,450 | |
| | HCMV IgG[2] | | | -4,314 | 3,556 | -0,159 | -1,213 | 0,230 | | | -21,887 | 47,589 | -0,059 | -0,460 | 0,647 | |
| | RRMS[3] | | | -2,470 | 3,031 | -0,104 | -0,815 | 0,418 | | | **-86,825** | **39,577** | **-0,276** | **-2,194** | **0,032** | |
| **TemRA CD8$^+$ T cells** | | F(4,58)= 4,449 | **0,003** | | | | | | F(4,56)= 2,627 | **0,044** | | | | | | 3H |
| | Sex[1] | | | -0,716 | 2,554 | -0,033 | -0,280 | 0,780 | | | 5,595 | 35,129 | 0,020 | 0,159 | 0,874 | |
| | Age | | | 0,212 | 0,145 | 0,178 | 1,460 | 0,150 | | | 0,344 | 1,973 | 0,023 | 0,174 | 0,862 | |
| | HCMV IgG[2] | | | **6,072** | **2,870** | **0,257** | **2,116** | **0,039** | | | 42,496 | 40,086 | 0,135 | 1,060 | 0,294 | |
| | RRMS[3] | | | **-5,980** | **2,447** | **-0,290** | **-2,444** | **0,018** | | | **-91,717** | **33,338** | **-0,345** | **-2,751** | **0,008** | |

[1] Reference is "Female"; [2] Reference is "IgG negative"; [3] Reference is "Healthy"

**HCMV,** Human Cytomegalovirus; **CM**, Central Memory; **EM**, Effector memory; **TemRA**, Terminally differentiated CD45RA-expressing memory cells;

**eTable 6**. Multiple linear regression models to evaluate the impact of sex (female vs. male), age, HCMV IgG seroprevalence and disease status (healthy vs. RRMS) on chemokines receptors expressing CD8$^+$ T cell subsets.

| Dependent Variable | Independent Variable | Percentage of cells: Model | *p*-value | B | SE | *β* | *t* | *p*-value | Absolute number of cells: Model | *p*-value | B | SE | *β* | *t* | *p*-value | Fig |
|---|---|---|---|---|---|---|---|---|---|---|---|---|---|---|---|---|
| **CCR4$^+$ Memory CD8$^+$ T cells** | | F(4,58)= 2,281 | 0,071 | | | | | | F(4,56)= 3,445 | **0,014** | | | | | | e2A |
| | Sex[1] | | | 0,307 | 1,516 | 0,026 | 0,203 | 0,840 | | | 9,077 | 7,095 | 0,159 | 1,279 | 0,206 | |
| | Age | | | **0,229** | **0,086** | **0,344** | **2,656** | **0,010** | | | **1,126** | **0,398** | **0,357** | **2,825** | **0,007** | |
| | HCMV IgG[2] | | | 0,312 | 1,704 | 0,024 | 0,183 | 0,855 | | | 2,927 | 8,096 | 0,045 | 0,362 | 0,719 | |
| | RRMS[3] | | | 1,118 | 1,452 | 0,097 | 0,770 | 0,445 | | | -4,544 | 6,733 | -0,083 | -0,675 | 0,503 | |
| **CCR6$^+$ Memory CD8$^+$ T cells** | | F(4,58)= 4,323 | **0,004** | | | | | | F(4,56)= 2,462 | 0,056 | | | | | | |
| | Sex[1] | | | **-6,404** | **2,064** | **-0,372** | **-3,103** | **0,003** | | | **-26,702** | **12,986** | **-0,263** | **-2,056** | **0,044** | |
| | Age | | | 0,148 | 0,117 | 0,154 | 1,262 | 0,212 | | | 1,022 | 0,729 | 0,182 | 1,401 | 0,167 | |
| | HCMV IgG[2] | | | **-6,556** | **2,319** | **-0,344** | **-2,827** | **0,006** | | | -24,516 | 14,818 | -0,212 | -1,654 | 0,104 | |
| | RRMS[3] | | | 0,324 | 1,977 | 0,020 | 0,164 | 0,870 | | | **-24,982** | **12,323** | **-0,256** | **-2,027** | **0,047** | |
| **CCR10$^+$ Memory CD8$^+$ T cells** | | F(4,58)= 4,690 | **0,002** | | | | | | F(4,56)= 3,582 | **0,011** | | | | | | |
| | Sex[1] | | | -0,592 | 0,555 | -0,127 | -1,067 | 0,290 | | | -0,364 | 3,023 | -0,015 | -0,121 | 0,904 | |
| | Age | | | **0,132** | **0,032** | **0,508** | **4,192** | **0,000** | | | **0,615** | **0,170** | **0,456** | **3,623** | **0,001** | |
| | HCMV IgG[2] | | | -0,679 | 0,623 | -0,131 | -1,089 | 0,281 | | | -1,015 | 3,450 | -0,036 | -0,294 | 0,770 | |
| | RRMS[3] | | | 0,406 | 0,531 | 0,090 | 0,765 | 0,447 | | | -1,893 | 2,869 | -0,081 | -0,660 | 0,512 | |
| **CxCR3$^+$ Memory CD8$^+$ T cells** | | F(4,58)= 1,548 | 0,200 | | | | | | F(4,56)= 0,924 | 0,456 | | | | | | |
| | Sex[1] | | | **-2,505** | **1,198** | **-0,271** | **-2,092** | **0,041** | | | -8,692 | 5,296 | -0,221 | -1,641 | 0,106 | |
| | Age | | | -0,007 | 0,068 | -0,014 | -0,109 | 0,913 | | | 0,267 | 0,297 | 0,122 | 0,896 | 0,374 | |
| | HCMV IgG[2] | | | -1,186 | 1,346 | -0,116 | -0,882 | 0,382 | | | -2,130 | 6,043 | -0,047 | -0,352 | 0,726 | |
| | RRMS[3] | | | 0,539 | 1,147 | 0,061 | 0,470 | 0,640 | | | -4,772 | 5,026 | -0,126 | -0,949 | 0,346 | |
| **CCR4$^+$ CM CD8$^+$ T cells** | | F(4,58)= 2,356 | 0,064 | | | | | | F(4,56)= 2,060 | 0,098 | | | | | | e2B |
| | Sex[1] | | | 2,700 | 2,843 | 0,120 | 0,950 | 0,346 | | | 3,735 | 3,462 | 0,140 | 1,079 | 0,285 | |
| | Age | | | **0,371** | **0,162** | **0,296** | **2,292** | **0,026** | | | **0,400** | **0,194** | **0,271** | **2,055** | **0,045** | |
| | HCMV IgG[2] | | | 1,586 | 3,195 | 0,064 | 0,496 | 0,621 | | | 2,630 | 3,950 | 0,086 | 0,666 | 0,508 | |
| | RRMS[3] | | | -1,534 | 2,723 | -0,071 | -0,563 | 0,575 | | | 0,047 | 3,285 | 0,002 | 0,014 | 0,989 | |
| **CCR6$^+$ CM CD8$^+$ T cells** | | F(4,58)= 5,716 | **0,001** | | | | | | F(4,56)= 3,878 | **0,008** | | | | | | |
| | Sex[1] | | | **-2,515** | **1,056** | **-0,275** | **-2,381** | **0,021** | | | **-1,998** | **0,810** | **-0,303** | **-2,467** | **0,017** | |
| | Age | | | -0,074 | 0,060 | -0,145 | -1,226 | 0,225 | | | 0,017 | 0,045 | 0,046 | 0,366 | 0,716 | |
| | HCMV IgG[2] | | | **-3,783** | **1,187** | **-0,374** | **-3,186** | **0,002** | | | **-2,515** | **0,924** | **-0,334** | **-2,721** | **0,009** | |
| | RRMS[3] | | | 0,243 | 1,012 | 0,028 | 0,240 | 0,811 | | | -1,510 | 0,769 | -0,238 | -1,965 | 0,054 | |
| **CCR10$^+$ CM CD8$^+$ T cells** | | F(4,58)= 3,178 | **0,020** | | | | | | F(4,56)= 2,242 | 0,076 | | | | | | |
| | Sex[1] | | | -1,512 | 1,050 | -0,178 | -1,440 | 0,155 | | | -0,462 | 1,107 | -0,054 | -0,417 | 0,678 | |
| | Age | | | **0,210** | **0,060** | **0,443** | **3,508** | **0,001** | | | **0,183** | **0,062** | **0,385** | **2,941** | **0,005** | |
| | HCMV IgG[2] | | | -0,744 | 1,180 | -0,079 | -0,630 | 0,531 | | | -0,178 | 1,263 | -0,018 | -0,141 | 0,888 | |
| | RRMS[3] | | | -0,521 | 1,006 | -0,064 | -0,518 | 0,607 | | | -0,121 | 1,050 | -0,015 | -0,115 | 0,909 | |
| **CxCR3$^+$ CM CD8$^+$ T cells** | | F(4,58)= 1,235 | 0,306 | | | | | | F(4,56)= 0,718 | 0,583 | | | | | | |
| | Sex[1] | | | -6,481 | 3,817 | -0,223 | -1,698 | 0,095 | | | -4,536 | 3,414 | -0,180 | -1,329 | 0,189 | |
| | Age | | | 0,016 | 0,217 | 0,010 | 0,072 | 0,943 | | | 0,256 | 0,192 | 0,183 | 1,333 | 0,188 | |
| | HCMV IgG[2] | | | -2,531 | 4,290 | -0,079 | -0,590 | 0,557 | | | -1,075 | 3,895 | -0,037 | -0,276 | 0,784 | |
| | RRMS[3] | | | 3,904 | 3,657 | 0,139 | 1,068 | 0,290 | | | -1,246 | 3,240 | -0,051 | -0,385 | 0,702 | |
| **CCR4$^+$ EM CD8$^+$ T cells** | | F(4,58)= 5,047 | **0,001** | | | | | | F(4,56)= 4,927 | **0,002** | | | | | | e2C |
| | Sex[1] | | | 0,020 | 1,285 | 0,002 | 0,015 | 0,988 | | | 4,751 | 3,554 | 0,160 | 1,337 | 0,187 | |
| | Age | | | **0,303** | **0,073** | **0,497** | **4,143** | **0,000** | | | **0,700** | **0,200** | **0,425** | **3,507** | **0,001** | |
| | HCMV IgG[2] | | | 0,522 | 1,444 | 0,043 | 0,361 | 0,719 | | | -0,384 | 4,056 | -0,011 | -0,095 | 0,925 | |
| | RRMS[3] | | | 0,052 | 1,231 | 0,005 | 0,042 | 0,967 | | | -4,322 | 3,373 | -0,151 | -1,281 | 0,205 | |
| **CCR6$^+$ EM CD8$^+$ T cells** | | F(4,58)= 6,089 | **0,000** | | | | | | F(4,56)= 3,400 | **0,015** | | | | | | |
| | Sex[1] | | | **-8,823** | **2,810** | **-0,360** | **-3,140** | **0,003** | | | **-22,271** | **9,472** | **-0,293** | **-2,351** | **0,022** | |
| | Age | | | 0,176 | 0,160 | 0,129 | 1,100 | 0,276 | | | 0,604 | 0,532 | 0,144 | 1,135 | 0,261 | |
| | HCMV IgG[2] | | | **-10,056** | **3,158** | **-0,371** | **-3,184** | **0,002** | | | **-30,853** | **10,808** | **-0,355** | **-2,855** | **0,006** | |
| | RRMS[3] | | | 3,989 | 2,692 | 0,169 | 1,482 | 0,144 | | | 0,667 | 8,989 | 0,009 | 0,074 | 0,941 | |

[1] Reference is "Female"; [2] Reference is "IgG negative"; [3] Reference is "Healthy"

**HCMV**, Human Cytomegalovirus; **CM**, Central Memory; **EM**, Efector Memory; **TemRA**, Terminally differentiated CD45RA-expressing memory cells;

**eTable 6 (cont.)**

| Dependent Variable | Independent Variable | Percentage of cells: Model | *p*-value | B | SE | *β* | *t* | *p*-value | Absolute number of cells: Model | *p*-value | B | SE | *β* | *t* | *p*-value | Fig |
|---|---|---|---|---|---|---|---|---|---|---|---|---|---|---|---|---|
| **CCR10[+] EM CD8[+] T cells** | | F(4,58)= 7,920 | **0,000** | | | | | | F(4,56)= 4,375 | **0,004** | | | | | | |
| | Sex[1] | | | -0,740 | 0,550 | -0,148 | -1,346 | 0,184 | | | -0,614 | 1,841 | -0,040 | -0,333 | 0,740 | |
| | Age | | | **0,174** | **0,031** | **0,623** | **5,555** | **0,000** | | | **0,423** | **0,103** | **0,504** | **4,094** | **0,000** | |
| | HCMV IgG[2] | | | -0,402 | 0,618 | -0,073 | -0,651 | 0,517 | | | -1,268 | 2,101 | -0,073 | -0,603 | 0,549 | |
| | RRMS[3] | | | 0,273 | 0,527 | 0,057 | 0,518 | 0,606 | | | -0,866 | 1,747 | -0,059 | -0,495 | 0,622 | |
| **CxCR3[+] EM CD8[+] T cells** | | F(4,58)= 0,801 | 0,530 | | | | | | F(4,56)= 0,823 | 0,516 | | | | | | |
| | Sex[1] | | | -1,401 | 0,845 | -0,220 | -1,658 | 0,103 | | | -2,390 | 2,026 | -0,159 | -1,180 | 0,243 | |
| | Age | | | 0,046 | 0,048 | 0,130 | 0,957 | 0,343 | | | 0,134 | 0,114 | 0,162 | 1,181 | 0,243 | |
| | HCMV IgG[2] | | | 0,128 | 0,950 | 0,018 | 0,135 | 0,893 | | | -0,208 | 2,312 | -0,012 | -0,090 | 0,928 | |
| | RRMS[3] | | | -0,142 | 0,810 | -0,023 | -0,175 | 0,862 | | | -2,067 | 1,923 | -0,143 | -1,075 | 0,287 | |
| **CCR4[+] TemRA CD8[+] T cells** | | F(4,58)= 0,377 | 0,824 | | | | | | F(4,56)= 1,042 | 0,394 | | | | | | e2D |
| | Sex[1] | | | -0,144 | 0,532 | -0,037 | -0,271 | 0,787 | | | 0,294 | 0,491 | 0,080 | 0,599 | 0,552 | |
| | Age | | | 0,006 | 0,030 | 0,028 | 0,206 | 0,837 | | | 0,036 | 0,028 | 0,176 | 1,291 | 0,202 | |
| | HCMV IgG[2] | | | 0,297 | 0,597 | 0,068 | 0,497 | 0,621 | | | 0,244 | 0,560 | 0,058 | 0,436 | 0,665 | |
| | RRMS[3] | | | 0,582 | 0,509 | 0,153 | 1,144 | 0,257 | | | -0,442 | 0,466 | -0,125 | -0,949 | 0,347 | |
| **CCR6[+] TemRA CD8[+] T cells** | | F(4,58)= 3,949 | **0,007** | | | | | | F(4,56)= 3,788 | **0,009** | | | | | | |
| | Sex[1] | | | -1,563 | 1,877 | -0,101 | -0,832 | 0,409 | | | -2,595 | 7,543 | -0,042 | -0,344 | 0,732 | |
| | Age | | | **0,294** | **0,107** | **0,340** | **2,749** | **0,008** | | | 0,385 | 0,424 | 0,114 | 0,908 | 0,368 | |
| | HCMV IgG[2] | | | -0,595 | 2,110 | -0,035 | -0,282 | 0,779 | | | 8,152 | 8,608 | 0,117 | 0,947 | 0,348 | |
| | RRMS[3] | | | **-5,086** | **1,799** | **-0,340** | **-2,828** | **0,006** | | | **-24,067** | **7,159** | **-0,408** | **-3,362** | **0,001** | |
| **CCR10[+] TemRA CD8[+] T cells** | | F(4,58)= 0,410 | 0,801 | | | | | | F(4,56)= 0,673 | 0,613 | | | | | | |
| | Sex[1] | | | 0,298 | 0,367 | 0,109 | 0,812 | 0,420 | | | 0,762 | 0,848 | 0,122 | 0,899 | 0,372 | |
| | Age | | | -0,016 | 0,021 | -0,108 | -0,785 | 0,436 | | | 0,010 | 0,048 | 0,030 | 0,217 | 0,829 | |
| | HCMV IgG[2] | | | -0,234 | 0,413 | -0,077 | -0,566 | 0,573 | | | 0,417 | 0,967 | 0,058 | 0,431 | 0,668 | |
| | RRMS[3] | | | 0,034 | 0,352 | 0,013 | 0,096 | 0,924 | | | -0,867 | 0,804 | -0,144 | -1,078 | 0,286 | |
| **CxCR3[+] TemRA CD8[+] T cells** | | F(4,58)= 4,290 | **0,004** | | | | | | F(4,56)= 1,858 | 0,131 | | | | | | |
| | Sex[1] | | | -0,931 | 0,471 | -0,237 | -1,977 | 0,053 | | | **-0,949** | **0,408** | **-0,303** | **-2,327** | **0,024** | |
| | Age | | | -0,041 | 0,027 | -0,187 | -1,530 | 0,131 | | | -0,006 | 0,023 | -0,034 | -0,255 | 0,800 | |
| | HCMV IgG[2] | | | **-1,154** | **0,529** | **-0,266** | **-2,180** | **0,033** | | | -0,359 | 0,465 | -0,101 | -0,771 | 0,444 | |
| | RRMS[3] | | | 0,398 | 0,451 | 0,105 | 0,883 | 0,381 | | | -0,325 | 0,387 | -0,108 | -0,839 | 0,405 | |

[1] Reference is "Female"; [2] Reference is "IgG negative"; [3] Reference is "Healthy"

**HCMV**, Human Cytomegalovirus; **CM**, Central Memory; **EM**, Efector Memory; **TemRA**, Terminally differentiated CD45RA-expressing memory cells;

**eTable 7.** Multiple linear regression models to evaluate the impact of sex (female *vs.* male), age, HCMV IgG seroprevalence and disease status (healthy *vs.* RRMS) on regulatory T cells (Treg) subsets.

| Dependent Variable | Independent Variable | Percentage of cells: Model | *p*-value | B | SE | *β* | *t* | *p*-value | Absolute number of cells: Model | *p*-value | B | SE | *β* | *t* | *p*-value | Fig. |
|---|---|---|---|---|---|---|---|---|---|---|---|---|---|---|---|---|
| **Treg** | | F(4,58)= 1,791 | 0,143 | | | | | | F(4,56)= 0,356 | 0,839 | | | | | | 4A |
| | Sex[1] | | | 0,898 | 0,471 | 0,245 | 1,906 | 0,062 | | | -1,526 | 8,599 | -0,024 | -0,178 | 0,860 | |
| | Age | | | -0,017 | 0,027 | -0,086 | -0,652 | 0,517 | | | 0,351 | 0,483 | 0,101 | 0,727 | 0,470 | |
| | HCMV IgG[2] | | | -0,598 | 0,530 | -0,148 | -1,129 | 0,264 | | | -8,837 | 9,812 | -0,123 | -0,901 | 0,372 | |
| | RRMS[3] | | | 0,451 | 0,452 | 0,128 | 0,999 | 0,322 | | | 2,597 | 8,160 | 0,043 | 0,318 | 0,751 | |
| **Naive/Activated Tregs[4]** | | F(4,58)= 4,638 | **0,003** | | | | | | | | | | | | | 4C |
| | Sex[1] | | | 0,116 | 0,090 | 0,154 | 1,295 | 0,200 | | | | | | | | |
| | Age | | | **-0,019** | **0,005** | **-0,445** | **-3,663** | **0,001** | | | | | | | | |
| | HCMV IgG[2] | | | -0,015 | 0,101 | -0,018 | -0,150 | 0,881 | | | | | | | | |
| | RRMS[3] | | | 0,166 | 0,086 | 0,227 | 1,923 | 0,059 | | | | | | | | |
| **CD45RA$^{+}$HLA-DR$^{-}$ Tregs[4]** | | F(4,58)= 5,688 | **0,001** | | | | | | F(4,56)= 1,950 | 0,115 | | | | | | 4D |
| | Sex[1] | | | 4,905 | 2,938 | 0,193 | 1,670 | 0,100 | | | 3,945 | 3,407 | 0,151 | 1,158 | 0,252 | |
| | Age | | | **-0,666** | **0,167** | **-0,471** | **-3,986** | **0,000** | | | **-0,432** | **0,191** | **-0,299** | **-2,260** | **0,028** | |
| | HCMV IgG[2] | | | 3,348 | 3,301 | 0,119 | 1,014 | 0,315 | | | 0,034 | 3,888 | 0,001 | 0,009 | 0,993 | |
| | RRMS[3] | | | **7,635** | **2,814** | **0,312** | **2,713** | **0,009** | | | 4,917 | 3,234 | 0,195 | 1,520 | 0,134 | |
| **CD45RA$^{-}$HLA-DR$^{-}$ Tregs[4]** | | F(4,58)= 4,055 | **0,006** | | | | | | F(4,56)= 1,107 | 0,362 | | | | | | 4E |
| | Sex[1] | | | **-6,167** | **2,083** | **-0,357** | **-2,961** | **0,004** | | | -5,240 | 4,282 | -0,164 | -1,224 | 0,226 | |
| | Age | | | **0,291** | **0,118** | **0,302** | **2,452** | **0,017** | | | 0,440 | 0,240 | 0,249 | 1,830 | 0,073 | |
| | HCMV IgG[2] | | | 0,237 | 2,341 | 0,012 | 0,101 | 0,920 | | | -5,002 | 4,886 | -0,137 | -1,024 | 0,310 | |
| | RRMS[3] | | | -3,839 | 1,996 | -0,231 | -1,924 | 0,059 | | | -0,746 | 4,063 | -0,024 | -0,184 | 0,855 | |
| **CD45RA$^{-}$HLA-DR$^{+}$ Tregs[4]** | | F(4,58)= 3,577 | **0,011** | | | | | | F(4,56)= 1,969 | 0,112 | | | | | | 4F |
| | Sex[1] | | | 1,128 | 2,219 | 0,062 | 0,508 | 0,613 | | | -0,349 | 2,448 | -0,019 | -0,143 | 0,887 | |
| | Age | | | **0,383** | **0,126** | **0,379** | **3,033** | **0,004** | | | **0,350** | **0,137** | **0,336** | **2,542** | **0,014** | |
| | HCMV IgG[2] | | | -3,692 | 2,494 | -0,184 | -1,480 | 0,144 | | | -3,934 | 2,793 | -0,183 | -1,408 | 0,165 | |
| | RRMS[3] | | | -3,995 | 2,126 | -0,228 | -1,879 | 0,065 | | | -1,749 | 2,323 | -0,096 | -0,753 | 0,455 | |
| **CD39$^{+}$ Tregs[4]** | | F(4,58)= 2,737 | **0,037** | | | | | | F(4,56)= 2,315 | 0,068 | | | | | | 4G |
| | Sex[1] | | | -2,944 | 3,739 | -0,099 | -0,787 | 0,434 | | | -3,489 | 3,757 | -0,119 | -0,929 | 0,357 | |
| | Age | | | **0,513** | **0,213** | **0,308** | **2,412** | **0,019** | | | **0,524** | **0,211** | **0,324** | **2,482** | **0,016** | |
| | HCMV IgG[2] | | | -5,834 | 4,201 | -0,177 | -1,389 | 0,170 | | | -7,953 | 4,287 | -0,238 | -1,855 | 0,069 | |
| | RRMS[3] | | | **-8,448** | **3,582** | **-0,294** | **-2,359** | **0,022** | | | -5,064 | 3,566 | -0,180 | -1,420 | 0,161 | |
| **CD73$^{+}$ Tregs[4]** | | F(4,58)= 0,500 | 0,736 | | | | | | F(4,56)= 1,303 | 0,280 | | | | | | 4H |
| | Sex[1] | | | -0,752 | 0,696 | -0,145 | -1,080 | 0,284 | | | -0,928 | 0,537 | -0,229 | -1,727 | 0,090 | |
| | Age | | | -0,015 | 0,040 | -0,053 | -0,389 | 0,699 | | | 0,013 | 0,030 | 0,059 | 0,441 | 0,661 | |
| | HCMV IgG[2] | | | -0,373 | 0,782 | -0,065 | -0,478 | 0,635 | | | -0,936 | 0,613 | -0,203 | -1,527 | 0,132 | |
| | RRMS[3] | | | -0,147 | 0,666 | -0,029 | -0,220 | 0,827 | | | -0,281 | 0,510 | -0,072 | -0,551 | 0,584 | |
| **GARP$^{+}$ Tregs[4]** | | F(4,58)= 2,285 | 0,071 | | | | | | F(4,56)= 1,560 | 0,198 | | | | | | 4I |
| | Sex[1] | | | 1,158 | 1,223 | 0,120 | 0,947 | 0,348 | | | 1,294 | 1,428 | 0,119 | 0,906 | 0,369 | |
| | Age | | | -0,071 | 0,070 | -0,133 | -1,027 | 0,309 | | | -0,051 | 0,080 | -0,085 | -0,635 | 0,528 | |
| | HCMV IgG[2] | | | 2,305 | 1,374 | 0,216 | 1,678 | 0,099 | | | 1,782 | 1,629 | 0,144 | 1,094 | 0,279 | |
| | RRMS[3] | | | **3,126** | **1,171** | **0,337** | **2,669** | **0,010** | | | **3,066** | **1,355** | **0,294** | **2,263** | **0,028** | |

[1] Reference is "Female"; [2] Reference is "IgG negative"; [3] Reference is "Healthy"; [4] Evaluated only when parent population >500 events
**HCMV,** Human Cytomegalovirus; **Treg**, Regulatory T cells

**eTable 8.** Multiple linear regression models to evaluate the impact of sex (female *vs.* male), age, HCMV IgG seroprevalence and disease status (healthy *vs.* RRMS) on natural killer (NK) cells subsets.

| Dependent Variable | Independent Variable | Percentage of cells: Model | *p*-value | B | SE | *β* | *t* | *p*-value | Absolute number of cells: Model | *p*-value | B | SE | *β* | *t* | *p*-value | Fig |
|---|---|---|---|---|---|---|---|---|---|---|---|---|---|---|---|---|
| **NK cells** | | F(4,43)= 1,502 | 0,219 | | | | | | F(4,38)= 0,795 | 0,536 | | | | | | 5A |
| | Sex[1] | | | 4,084 | 2,404 | 0,258 | 1,698 | 0,097 | | | 116,994 | 124,290 | 0,157 | 0,941 | 0,352 | |
| | Age | | | 0,116 | 0,106 | 0,163 | 1,096 | 0,279 | | | 5,612 | 5,290 | 0,171 | 1,061 | 0,296 | |
| | HCMV IgG[2] | | | -1,925 | 2,200 | -0,133 | -0,875 | 0,387 | | | -47,098 | 113,728 | -0,068 | -0,414 | 0,681 | |
| | RRMS[3] | | | -0,343 | 1,977 | -0,027 | -0,173 | 0,863 | | | 32,989 | 102,687 | 0,054 | 0,321 | 0,750 | |
| **CD56$^{bright}$ NK cells** | | F(4,43)= 2,806 | **0,037** | | | | | | F(4,38)= 1,377 | 0,260 | | | | | | 5B |
| | Sex[1] | | | -1,984 | 1,629 | -0,176 | -1,218 | 0,230 | | | -6,902 | 4,811 | -0,233 | -1,435 | 0,160 | |
| | Age | | | **-0,165** | **0,072** | **-0,325** | **-2,303** | **0,026** | | | -0,189 | 0,205 | -0,144 | -0,922 | 0,362 | |
| | HCMV IgG[2] | | | 1,823 | 1,491 | 0,177 | 1,223 | 0,228 | | | 0,550 | 4,402 | 0,020 | 0,125 | 0,901 | |
| | RRMS[3] | | | 2,508 | 1,340 | 0,273 | 1,872 | 0,068 | | | 6,721 | 3,974 | 0,279 | 1,691 | 0,099 | |
| **CD56$^{dim}$CD57$^-$ NK cells** | | F(4,43)= 1,971 | 0,116 | | | | | | F(4,38)= 1,068 | 0,386 | | | | | | 5C |
| | Sex[1] | | | -2,713 | 4,467 | -0,091 | -0,607 | 0,547 | | | 18,965 | 59,938 | 0,052 | 0,316 | 0,753 | |
| | Age | | | 0,011 | 0,196 | 0,008 | 0,057 | 0,954 | | | 2,834 | 2,551 | 0,176 | 1,111 | 0,274 | |
| | HCMV IgG[2] | | | **-10,633** | **4,087** | **-0,388** | **-2,601** | **0,013** | | | -78,910 | 54,845 | -0,233 | -1,439 | 0,158 | |
| | RRMS[3] | | | 0,926 | 3,673 | 0,038 | 0,252 | 0,802 | | | 23,535 | 49,520 | 0,080 | 0,475 | 0,637 | |
| **CD56$^{dim}$CD57$^+$ NK cells** | | F(4,43)= 1,856 | 0,136 | | | | | | F(4,38)= 1,227 | 0,316 | | | | | | 5D |
| | Sex[1] | | | 4,802 | 4,970 | 0,145 | 0,966 | 0,339 | | | 104,821 | 67,323 | 0,255 | 1,557 | 0,128 | |
| | Age | | | 0,159 | 0,218 | 0,107 | 0,728 | 0,470 | | | 2,968 | 2,866 | 0,163 | 1,036 | 0,307 | |
| | HCMV IgG[2] | | | 8,620 | 4,548 | 0,284 | 1,895 | 0,065 | | | 30,776 | 61,602 | 0,080 | 0,500 | 0,620 | |
| | RRMS[3] | | | -3,599 | 4,087 | -0,133 | -0,881 | 0,383 | | | 2,247 | 55,621 | 0,007 | 0,040 | 0,968 | |
| **KLRG1$^+$ CD56$^{bright}$ NK cells[4]** | | F(4,33)= 3,949 | **0,010** | | | | | | | | | | | | | 5E |
| | Sex[1] | | | **8,124** | **2,935** | **0,418** | **2,768** | **0,009** | | | | | | | | |
| | Age | | | -0,054 | 0,132 | -0,060 | -0,410 | 0,684 | | | | | | | | |
| | HCMV IgG[2] | | | 0,137 | 2,707 | 0,008 | 0,051 | 0,960 | | | | | | | | |
| | RRMS[3] | | | **5,128** | **2,429** | **0,322** | **2,111** | **0,042** | | | | | | | | |
| **KLRG1$^+$ CD56$^{dim}$CD57$^-$ NK cells[4]** | | F(4,41)= 0,031 | 0,998 | | | | | | | | | | | | | |
| | Sex[1] | | | -1,914 | 6,880 | -0,046 | -0,278 | 0,782 | | | | | | | | |
| | Age | | | 0,066 | 0,299 | 0,036 | 0,220 | 0,827 | | | | | | | | |
| | HCMV IgG[2] | | | -0,319 | 6,390 | -0,008 | -0,050 | 0,960 | | | | | | | | |
| | RRMS[3] | | | -0,415 | 5,585 | -0,013 | -0,074 | 0,941 | | | | | | | | |
| **KLRG1$^+$ CD56$^{dim}$CD57$^+$ NK cells[4]** | | F(4,41)= 0,678 | 0,611 | | | | | | | | | | | | | |
| | Sex[1] | | | -3,575 | 8,479 | -0,068 | -0,422 | 0,675 | | | | | | | | |
| | Age | | | 0,206 | 0,368 | 0,088 | 0,558 | 0,580 | | | | | | | | |
| | HCMV IgG[2] | | | 7,798 | 7,875 | 0,158 | 0,990 | 0,328 | | | | | | | | |
| | RRMS[3] | | | -3,915 | 6,883 | -0,093 | -0,569 | 0,573 | | | | | | | | |
| **NKG2A$^+$ CD56$^{bright}$ NK cells[4]** | | F(4,33)= 1,086 | 0,380 | | | | | | | | | | | | | 5F |
| | Sex[1] | | | -1,066 | 1,279 | -0,144 | -0,834 | 0,410 | | | | | | | | |
| | Age | | | -0,037 | 0,058 | -0,107 | -0,636 | 0,529 | | | | | | | | |
| | HCMV IgG[2] | | | -0,956 | 1,180 | -0,139 | -0,811 | 0,423 | | | | | | | | |
| | RRMS[3] | | | 1,439 | 1,059 | 0,237 | 1,359 | 0,183 | | | | | | | | |
| **NKG2A$^+$ CD56$^{dim}$CD57$^-$ NK cells[4]** | | F(4,41)= 1,387 | 0,255 | | | | | | | | | | | | | |
| | Sex[1] | | | -1,148 | 4,688 | -0,038 | -0,245 | 0,808 | | | | | | | | |
| | Age | | | -0,128 | 0,204 | -0,095 | -0,627 | 0,534 | | | | | | | | |
| | HCMV IgG[2] | | | -4,799 | 4,354 | -0,171 | -1,102 | 0,277 | | | | | | | | |
| | RRMS[3] | | | 5,508 | 3,805 | 0,229 | 1,447 | 0,155 | | | | | | | | |

[1] Reference is "Female"; [2] Reference is "IgG negative"; [3] Reference is "Healthy"; [4] Evaluated only when parent population >500 events
**HCMV**, Human Cytomegalovirus; **NK**, Natural Killer.

**eTable 9.** Multiple linear regression models to evaluate the impact of sex (female *vs.* male), age, HCMV IgG seroprevalence and disease status (healthy *vs.* RRMS) on NKT cells.

| Dependent Variable | Independent Variable | Percentage of cells: Model | *p*-value | B | SE | *β* | *t* | *p*-value | Absolute number of cells: Model | *p*-value | B | SE | *β* | *t* | *p*-value | Fig. |
|---|---|---|---|---|---|---|---|---|---|---|---|---|---|---|---|---|
| **NKT cells** | | F(4,43)= 1,485 | 0,224 | | | | | | F(4,38)= 0,746 | 0,576 | | | | | | e3A |
| | Sex[1] | | | 0,091 | 0,831 | 0,017 | 0,109 | 0,914 | | | 32,201 | 36,008 | 0,150 | 0,894 | 0,377 | |
| | Age | | | -0,009 | 0,037 | -0,036 | -0,241 | 0,811 | | | -0,040 | 1,533 | -0,004 | -0,026 | 0,979 | |
| | HCMV IgG[2] | | | **1,799** | **0,761** | **0,360** | **2,364** | **0,023** | | | 48,190 | 32,948 | 0,240 | 1,463 | 0,152 | |
| | RRMS[3] | | | 0,081 | 0,684 | 0,018 | 0,118 | 0,907 | | | 1,196 | 29,749 | 0,007 | 0,040 | 0,968 | |
| **KLRG1⁺ NKT cells** | | F(4,41)= 0,862 | 0,495 | | | | | | | | | | | | | e3B |
| | Sex[1] | | | 1,805 | 3,147 | 0,091 | 0,574 | 0,569 | | | | | | | | |
| | Age | | | -0,097 | 0,137 | -0,111 | -0,713 | 0,480 | | | | | | | | |
| | HCMV IgG[2] | | | -3,842 | 2,923 | -0,209 | -1,314 | 0,196 | | | | | | | | |
| | RRMS[3] | | | -3,543 | 2,554 | -0,225 | -1,387 | 0,173 | | | | | | | | |
| **NKG2A⁺ NKT cells** | | F(4,41)= 4,127 | **0,007** | | | | | | | | | | | | | e3C |
| | Sex[1] | | | **-10,597** | **5,153** | **-0,287** | **-2,056** | **0,046** | | | | | | | | |
| | Age | | | 0,103 | 0,224 | 0,063 | 0,462 | 0,647 | | | | | | | | |
| | HCMV IgG[2] | | | **-17,222** | **4,786** | **-0,502** | **-3,598** | **0,001** | | | | | | | | |
| | RRMS[3] | | | -1,336 | 4,183 | -0,045 | -0,319 | 0,751 | | | | | | | | |
| **KIR2DL2/3⁺ NKT cells** | | F(4,41)= 1,630 | 0,185 | | | | | | | | | | | | | e3D |
| | Sex[1] | | | -1,691 | 3,647 | -0,071 | -0,464 | 0,645 | | | | | | | | |
| | Age | | | -0,069 | 0,158 | -0,065 | -0,434 | 0,666 | | | | | | | | |
| | HCMV IgG[2] | | | 4,568 | 3,388 | 0,207 | 1,348 | 0,185 | | | | | | | | |
| | RRMS[3] | | | -4,190 | 2,961 | -0,222 | -1,415 | 0,165 | | | | | | | | |
| **KIR3DL1⁺ NKT cells** | | F(4,41)= 2,539 | 0,054 | | | | | | | | | | | | | e3E |
| | Sex[1] | | | -0,637 | 2,969 | -0,032 | -0,215 | 0,831 | | | | | | | | |
| | Age | | | -0,194 | 0,129 | -0,218 | -1,502 | 0,141 | | | | | | | | |
| | HCMV IgG[2] | | | **6,073** | **2,757** | **0,326** | **2,203** | **0,033** | | | | | | | | |
| | RRMS[3] | | | -2,520 | 2,410 | -0,158 | -1,046 | 0,302 | | | | | | | | |
| **NKp30⁺ NKT cells** | | F(4,41)= 2,762 | **0,040** | | | | | | | | | | | | | e3F |
| | Sex[1] | | | -3,772 | 3,406 | -0,163 | -1,107 | 0,275 | | | | | | | | |
| | Age | | | -0,281 | 0,146 | -0,278 | -1,922 | 0,062 | | | | | | | | |
| | HCMV IgG[2] | | | -4,633 | 3,032 | -0,221 | -1,528 | 0,134 | | | | | | | | |
| | RRMS[3] | | | 2,596 | 2,707 | 0,141 | 0,959 | 0,343 | | | | | | | | |
| **NKp44⁺ NKT cells** | | F(4,41)= 0,300 | 0,876 | | | | | | | | | | | | | e3G |
| | Sex[1] | | | 2,760 | 2,877 | 0,157 | 0,959 | 0,343 | | | | | | | | |
| | Age | | | -0,067 | 0,123 | -0,088 | -0,547 | 0,587 | | | | | | | | |
| | HCMV IgG[2] | | | 0,570 | 2,561 | 0,036 | 0,222 | 0,825 | | | | | | | | |
| | RRMS[3] | | | -1,292 | 2,287 | -0,092 | -0,565 | 0,575 | | | | | | | | |
| **NKp46⁺ NKT cells** | | F(4,41)= 1,267 | 0,298 | | | | | | | | | | | | | e3H |
| | Sex[1] | | | 0,166 | 0,786 | 0,033 | 0,212 | 0,834 | | | | | | | | |
| | Age | | | -0,043 | 0,034 | -0,196 | -1,272 | 0,211 | | | | | | | | |
| | HCMV IgG[2] | | | 0,760 | 0,699 | 0,167 | 1,087 | 0,283 | | | | | | | | |
| | RRMS[3] | | | 1,071 | 0,624 | 0,268 | 1,716 | 0,094 | | | | | | | | |

[1] Reference is "Female"; [2] Reference is "IgG negative"; [3] Reference is "Healthy"

**eTable 10.** Multiple linear regression models to evaluate the impact of time since last relapse on the several blood cell populations while controlling for corticoids administration to treat the relapse. The unstandardized residuals from a preliminary multiple linear regression were calculated, where the contribution of sex, age and HCMV IgG seroprevalence on the blood populations was evaluated. The residuals were used as dependent variable to evaluate the impact of time since last relapse and corticids on the blood cell populations. Only the blood populations for which the multiple linear regressions were significant ($p < 0.050$) for time from relapse are represented.

| Dependent Variable | Independent Variable | Percentage of cells | | | | | | | | Absolute number of cells | | | | | | | |
|---|---|---|---|---|---|---|---|---|---|---|---|---|---|---|---|---|---|
| | | Model | *p*-value | Effect Size ($R^2$) | B | SE | *β* | *t* | *p*-value | Model | *p*-value | Effect Size ($R^2$) | B | SE | *β* | *t* | *p*-value |
| **CD4⁺ T cells** | | | | | | | | | | F(2,24)= 7,857 | **0,002** | *0,396* | | | | | |
| | Last relapse (months) | | | | | | | | | | | | **26,244** | **6,620** | **0,636** | **3,964** | **0,001** |
| | Corticoids[1] | | | | | | | | | | | | 86,418 | 145,155 | 0,096 | 0,595 | 0,557 |
| **Naive CD4⁺ T cells** | | | | | | | | | | F(2,24)= 6,388 | **0,006** | *0,347* | | | | | |
| | Last relapse (months) | | | | | | | | | | | | **14,903** | **4,170** | **0,596** | **3,574** | **0,002** |
| | Corticoids[1] | | | | | | | | | | | | 46,422 | 91,422 | 0,085 | 0,508 | 0,616 |
| **Memory CD4⁺ T cells** | | | | | | | | | | F(2,24)= 4,337 | **0,025** | *0,265* | | | | | |
| | Last relapse (months) | | | | | | | | | | | | **11,340** | **3,851** | **0,521** | **2,945** | **0,007** |
| | Corticoids[1] | | | | | | | | | | | | 39,995 | 84,435 | 0,084 | 0,474 | 0,640 |
| **CM CD4⁺ T cells** | | | | | | | | | | F(2,24)= 5,568 | **0,010** | *0,317* | | | | | |
| | Last relapse (months) | | | | | | | | | | | | **9,218** | **2,887** | **0,545** | **3,192** | **0,004** |
| | Corticoids[1] | | | | | | | | | | | | -30,623 | 63,309 | -0,083 | -0,484 | 0,633 |
| **Th2** | | | | | | | | | | F(2,24)= 4,298 | **0,025** | *0,264* | | | | | |
| | Last relapse (months) | | | | | | | | | | | | **2,203** | **0,914** | **0,427** | **2,410** | **0,024** |
| | Corticoids[1] | | | | | | | | | | | | -25,869 | 20,042 | -0,229 | -1,291 | 0,209 |
| **CCR4⁺ Memory CD4⁺ T** | | | | | | | | | | F(2,24)= 3,657 | **0,041** | 0,234 | | | | | |
| | Last relapse (months) | | | | | | | | | | | | **4,213** | **1,804** | **0,422** | **2,336** | **0,028** |
| | Corticoids[1] | | | | | | | | | | | | -39,492 | 39,545 | -0,180 | -0,999 | 0,328 |
| **CCR4⁺ CM CD4⁺ T cells** | | | | | | | | | | F(2,24)= 5,737 | **0,009** | 0,323 | | | | | |
| | Last relapse (months) | | | | | | | | | | | | **3,002** | **1,162** | **0,439** | **2,583** | **0,016** |
| | Corticoids[1] | | | | | | | | | | | | -45,363 | 25,481 | -0,302 | -1,780 | 0,088 |
| **CCR10⁺ CM CD4⁺ T cells** | | | | | | | | | | F(2,24)= 4,070 | **0,030** | 0,253 | | | | | |
| | Last relapse (months) | | | | | | | | | | | | **1,163** | **0,439** | **0,473** | **2,650** | **0,014** |
| | Corticoids[1] | | | | | | | | | | | | -6,244 | 9,620 | -0,116 | -0,649 | 0,522 |
| **CCR6⁺ EM CD4⁺ T cells** | | F(2,24)= 3,556 | **0,043** | 0,215 | | | | | | | | | | | | | |
| | Last relapse (months) | | | | **-0,285** | **0,108** | **-0,462** | **-2,630** | **0,014** | | | | | | | | |
| | Corticoids[1] | | | | 0,098 | 2,307 | 0,007 | 0,043 | 0,966 | | | | | | | | |
| **CM CD8⁺ T cells** | | F(2,26)= 6,771 | **0,004** | 0,342 | | | | | | F(2,24)= 5,506 | **0,011** | 0,315 | | | | | |
| | Last relapse (months) | | | | **0,109** | **0,045** | **0,386** | **2,402** | **0,024** | | | | **1,634** | **0,496** | **0,562** | **3,291** | **0,003** |
| | Corticoids[1] | | | | **-2,316** | **0,965** | **-0,386** | **-2,399** | **0,024** | | | | 0,758 | 10,886 | 0,012 | 0,070 | 0,945 |
| **CCR4⁺ Memory CD8⁺ T** | | F(2,24)= 4,797 | **0,017** | 0,270 | | | | | | F(2,24)= 5,544 | **0,010** | 0,316 | | | | | |
| | Last relapse (months) | | | | **0,148** | **0,056** | **0,450** | **2,652** | **0,013** | | | | **1,019** | **0,320** | **0,544** | **3,183** | **0,004** |
| | Corticoids[1] | | | | -1,415 | 1,192 | -0,201 | -1,187 | 0,246 | | | | -3,442 | 7,021 | -0,084 | -0,490 | 0,628 |
| **CCR6⁺ Memory CD8⁺ T** | | F(2,24)= 6,732 | **0,004** | 0,341 | | | | | | | | | | | | | |
| | Last relapse (months) | | | | **-0,407** | **0,137** | **-0,479** | **-2,975** | **0,006** | | | | | | | | |
| | Corticoids[1] | | | | 4,908 | 2,920 | 0,271 | 1,681 | 0,105 | | | | | | | | |
| **CCR10⁺ Memory CD8⁺ T** | | F(2,24)= 3,631 | **0,041** | 0,218 | | | | | | F(2,24)= 5,179 | **0,013** | 0,301 | | | | | |
| | Last relapse (months) | | | | **0,073** | **0,028** | **0,459** | **2,620** | **0,014** | | | | **0,478** | **0,149** | **0,555** | **3,218** | **0,004** |
| | Corticoids[1] | | | | -0,140 | 0,596 | -0,041 | -0,235 | 0,816 | | | | 1,702 | 3,258 | 0,090 | 0,522 | 0,606 |
| **CCR4⁺ CM CD8⁺ T cells** | | F(2,24)= 6,948 | **0,004** | 0,348 | | | | | | F(2,24)= 8,335 | **0,002** | 0,410 | | | | | |
| | Last relapse (months) | | | | **0,360** | **0,107** | **0,537** | **3,354** | **0,002** | | | | **0,614** | **0,157** | **0,622** | **3,923** | **0,001** |
| | Corticoids[1] | | | | -2,535 | 2,286 | -0,178 | -1,109 | 0,278 | | | | -1,829 | 3,434 | -0,084 | -0,532 | 0,599 |
| **CCR6⁺ CM CD8⁺ T cells** | | F(2,24)= 5,098 | **0,014** | 0,282 | | | | | | | | | | | | | |
| | Last relapse (months) | | | | **-0,196** | **0,071** | **-0,461** | **-2,743** | **0,011** | | | | | | | | |
| | Corticoids[1] | | | | 1,838 | 1,522 | 0,203 | 1,208 | 0,238 | | | | | | | | |
| **CCR10⁺ CM CD8⁺ T cells** | | F(2,24)= 3,765 | **0,037** | 0,225 | | | | | | F(2,24)= 7,199 | **0,004** | 0,375 | | | | | |
| | Last relapse (months) | | | | **0,126** | **0,048** | **0,457** | **2,619** | **0,015** | | | | **0,200** | **0,053** | **0,616** | **3,774** | **0,001** |
| | Corticoids[1] | | | | -0,433 | 1,030 | -0,073 | -0,420 | 0,678 | | | | 0,206 | 1,164 | 0,029 | 0,177 | 0,861 |
| **CCR4⁺ EM CD8⁺ T cells** | | F(2,24)= 3,435 | **0,047** | 0,209 | | | | | | | | | | | | | |
| | Last relapse (months) | | | | **0,117** | **0,049** | **0,423** | **2,396** | **0,024** | | | | | | | | |
| | Corticoids[1] | | | | -0,727 | 1,045 | -0,123 | -0,695 | 0,493 | | | | | | | | |
| **CCR6⁺ EM CD8⁺ T cells** | | F(2,24)= 6,926 | **0,004** | 0,348 | | | | | | | | | | | | | |
| | Last relapse (months) | | | | **-0,567** | **0,177** | **-0,514** | **-3,210** | **0,004** | | | | | | | | |
| | Corticoids[1] | | | | 5,217 | 3,767 | 0,222 | 1,385 | 0,178 | | | | | | | | |

[1] Reference is "No history of corticoids"; **CM**, Central memory; **EM**, Effector Memory; **NK**, Natural Killer cells; **NKT**, Natural Killer cells; **TemRA**, Terminally differentiated CD45RA-expressing memory cells; **Treg**, Regulatory T cells

**eTable 10 (cont.)**

| Dependent Variable | Independent Variable | Percentage of cells: Model | *p*-value | Effect Size ($R^2$) | B | SE | *β* | *t* | *p*-value | Absolute number of cells: Model | *p*-value | Effect Size ($R^2$) | B | SE | *β* | *t* | *p*-value |
|---|---|---|---|---|---|---|---|---|---|---|---|---|---|---|---|---|---|
| **CCR10⁺ EM CD8⁺ T cells** | | F(2,24)= 4,509 | **0,021** | 0,258 | | | | | | F(2,24)= 4,391 | **0,024** | 0,268 | | | | | |
| | Last relapse (months) | | | | **0,091** | **0,030** | **0,511** | **2,993** | **0,006** | | | | **0,280** | **0,094** | **0,523** | **2,960** | **0,007** |
| | Corticoids[1] | | | | 0,128 | 0,650 | 0,034 | 0,197 | 0,845 | | | | 1,209 | 2,072 | 0,103 | 0,584 | 0,565 |
| **CCR10⁺ TemRA CD8⁺ T** | | F(2,24)= 4,721 | **0,018** | 0,266 | | | | | | | | | | | | | |
| | Last relapse (months) | | | | **-0,022** | **0,010** | **-0,373** | **-2,196** | **0,037** | | | | | | | | |
| | Corticoids[1] | | | | 0,377 | 0,210 | 0,305 | 1,798 | 0,084 | | | | | | | | |
| **Tregs** | | | | | | | | | | F(2,24)= 7,401 | **0,003** | *0,381* | | | | | |
| | Last relapse (months) | | | | | | | | | | | | **2,215** | **0,615** | **0,585** | **3,602** | **0,001** |
| | Corticoids[1] | | | | | | | | | | | | 25,290 | 13,483 | 0,305 | 1,876 | 0,073 |
| **CD45RA⁺HLA-DR⁻ Tregs** | | | | | | | | | | F(2,24)= 9,065 | **0,001** | 0,430 | | | | | |
| | Last relapse (months) | | | | | | | | | | | | **0,801** | **0,197** | **0,633** | **4,059** | **0,000** |
| | Corticoids[1] | | | | | | | | | | | | 8,119 | 4,326 | 0,292 | 1,877 | 0,073 |
| **CD45RA⁻HLA-DR⁻ Tregs** | | | | | | | | | | F(2,24)= 5,129 | **0,014** | 0,299 | | | | | |
| | Last relapse (months) | | | | | | | | | | | | **0,981** | **0,325** | **0,521** | **3,016** | **0,006** |
| | Corticoids[1] | | | | | | | | | | | | 10,819 | 7,131 | 0,262 | 1,517 | 0,142 |
| **CD45RA⁻HLA-DR⁺ Tregs** | | | | | | | | | | F(2,24)= 4,388 | **0,024** | 0,268 | | | | | |
| | Last relapse (months) | | | | | | | | | | | | **0,417** | **0,159** | **0,462** | **2,616** | **0,015** |
| | Corticoids[1] | | | | | | | | | | | | 6,170 | 3,496 | 0,312 | 1,765 | 0,090 |
| **CD39⁺ Tregs** | | | | | | | | | | F(2,24)= 4,345 | **0,025** | *0,266* | | | | | |
| | Last relapse (months) | | | | | | | | | | | | **0,685** | **0,239** | **0,507** | **2,865** | **0,009** |
| | Corticoids[1] | | | | | | | | | | | | 5,838 | 5,244 | 0,197 | 1,113 | 0,277 |
| **NK cells** | | | | | | | | | | F(2,18)= 3,018 | 0,074 | *0,251* | | | | | |
| | Last relapse (months) | | | | | | | | | | | | **25,902** | **11,729** | **0,456** | **2,208** | **0,040** |
| | Corticoids[1] | | | | | | | | | | | | 112,313 | 156,787 | 0,148 | 0,716 | 0,483 |
| **CD56^dim^CD57⁺ NK cells** | | | | | | | | | | F(2,18)= 4,928 | **0,020** | *0,354* | | | | | |
| | Last relapse (months) | | | | | | | | | | | | **16,404** | **5,607** | **0,561** | **2,925** | **0,009** |
| | Corticoids[1] | | | | | | | | | | | | 49,904 | 74,957 | 0,128 | 0,666 | 0,514 |
| **NKT cells** | | | | | | | | | | F(2,18)= 6,033 | **0,010** | *0,401* | | | | | |
| | Last relapse (months) | | | | | | | | | | | | **10,494** | **3,092** | **0,627** | **3,394** | **0,003** |
| | Corticoids[1] | | | | | | | | | | | | 8,161 | 41,332 | 0,036 | 0,197 | 0,846 |

**CM**, Central memory; **EM**, Effector Memory; **NK**, Natural Killer cells; **NKT**, Natural Killer cells; **TemRA**, Terminally differentiated CD45RA-expressing memory cells; **Th**, Helper T cells; **Treg**, Regulatory T cells

**eTable 11.** Linear regression models to evaluate the impact of the Multuple Sclerosis Severity Score (MSSS) on the several blood cell populations. The unstandardized residuals from a preliminary multiple linear regression were calculated where the contribution of sex, age and HCMV IgG seroprevalence on the blood populations was evaluated. The residuals were used as dependent variable to evaluate the impact of MSSS of the blood cell populations. Only the blood populations for which the linear regressions were significant (p <0.050) are represented.

| Dependent Variable | Independent Variable | Percentage of cells | | | | | | | Absolute number of cells | | | | | | |
|---|---|---|---|---|---|---|---|---|---|---|---|---|---|---|---|
| | | Model | Effect Size ($R^2$) | B | SE | *β* | *t* | *p-value* | Model | Effect Size ($R^2$) | B | SE | *β* | *t* | *p-value* |
| **sjTREC levels** | | | | | | | | | F(1,19)= 5,704 | 0,231 | | | | | |
| | MSSS | | | | | | | | | | **-69,351** | **29,039** | **-0,481** | **-2,388** | **0,027** |
| **Th17** | | F(1,28)= 4,607 | 0,141 | | | | | | F(1,26)= 4,430 | 0,146 | | | | | |
| | MSSS | | | **-0,113** | **0,053** | **-0,376** | **-2,146** | **0,041** | | | **-1,277** | **0,607** | **-0,382** | **-2,105** | **0,045** |
| **CCR6$^+$ Memory CD4$^+$ T cells** | | F(1,28)= 7,168 | 0,204 | | | | | | | | | | | | |
| | MSSS | | | **-0,984** | **0,367** | **-0,451** | **-2,677** | **0,012** | | | | | | | |
| **CCR6$^+$ CM CD4$^+$ T cells** | | F(1,28)= 7,380 | 0,209 | | | | | | | | | | | | |
| | MSSS | | | **-1,271** | **0,468** | **-0,457** | **-2,717** | **0,011** | | | | | | | |
| **CCR6$^+$ EM CD4$^+$ T cells** | | F(1,28)= 5,988 | 0,176 | | | | | | | | | | | | |
| | MSSS | | | **-0,928** | **0,379** | **-0,420** | **-2,447** | **0,021** | | | | | | | |
| **Naive/Activated Tregs** | | F(1,28)= 6,840 | 0,196 | | | | | | | | | | | | |
| | MSSS | | | **-0,039** | **0,015** | **-0,443** | **-2,615** | **0,014** | | | | | | | |
| **CD45RA$^+$HLA-DR$^-$ Tregs** | | F(1,28)= 6,783 | 0,195 | | | | | | | | | | | | |
| | MSSS | | | **-1,365** | **0,524** | **-0,442** | **-2,604** | **0,015** | | | | | | | |
| **GARP$^+$ Tregs** | | F(1,28)= 4,997 | 0,151 | | | | | | F(1,26)= 4,679 | 0,153 | | | | | |
| | MSSS | | | **0,722** | **0,323** | **0,389** | **2,235** | **0,034** | | | **0,875** | **0,404** | **0,391** | **2,163** | **0,040** |
| **KIR3DL1$^+$NKT cells** | | F(1,23)= 6,434 | 0,219 | | | | | | | | | | | | |
| | MSSS | | | **1,408** | **0,555** | **0,468** | **2,537** | **0,018** | | | | | | | |
| **KIR2DL2/3$^+$ NKT cells** | | F(1,23)= 5,818 | 0,202 | | | | | | | | | | | | |
| | MSSS | | | **0,661** | **0,274** | **0,449** | **2,412** | **0,024** | | | | | | | |

**CM,** Central memory; **EM,** Effector memory; **MSSS,** Multiple Sclerosis Severity Score; **NKT**, Natural Killer cells; **Th**, Helper T cells; **TREC**, T cell receptor excision circles; **Treg**, Regulatory T cells

**eTable 12.** Demographic and clinical characterization of the cohort.

| | **Newly diagnosed RRMS Patients (n = 30)** | **Healthy Controls (n = 33)** |
|---|---|---|
| Age (years). Mean [range] | 33.7 [19;54] | 33.7 [21;55][1] |
| Men. % (*n*) | 36.7 (11) | 36.4 (12)[2] |
| Age at MS onset (years). Median [range] | 31 [18;53] | na |
| Time since MS diagnosis (months). Median [range] | 1 [0;17] | na |
| Time from last relapse (months). Median [range] | 7 [0;49] | na |
| MSSS. Median [range] | 2.4 [0.7;9.4] | na |
| anti-HCMV $IgG^{+}$. % (*n*) | 63.3 (19) | 84.8 (28)[3] |

[1] $t_{61}$ = 0.000; $p > 0.9999$.

[2] $\chi^2$ (df=1) = 0.001; $p = 0.980$.

[3] $\chi^2$ (df=1) = 3.839; $p = 0.050$

**HCMV**, Human Cytomegalovirus; **MSSS**, Multiple Sclerosis Severity Score; **na**, not applicable

**eTable 13.** Panels of anti-human antibodies used for blood cells phenotypical characterization.

| Panel | Target | Clone | Fluorochrome | Excitation laser | Detector | Dilution[1] | Company |
|---|---|---|---|---|---|---|---|
| **Recent Thymic Emigrants T cell homeostasis** (~1 million PBMCs) | CD19 | HIB19 | FITC | 488 | 530/30 | 1:150 | BioLegend |
| | CD3 | OKT3 | PE | 488 | 575/26 | 1:200 | BioLegend |
| | 7AAD | na | na | 488 | 695/40 | 1:40 | BioLegend |
| | CD31 | WM59 | PE-Cy7 | 488 | 780/60 | 1:80 | BioLegend |
| | CD8 | RPA-T8 | APC | 633 | 660/20 | 1:150 | BioLegend |
| | CD4 | RPA-T4 | APC-Cy7 | 633 | 780/60 | 1:80 | BioLegend |
| | CD45RA | HI100 | Pacific Blue | 407 | 450/50 | 1:300 | BioLegend |
| | CD45 | HI30 | BV510 | 407 | 525/50 | 1:80 | BioLegend |
| | CD45RO | UCHL1 | BV650 | 407 | 660/20 | 1:300 | BioLegend |
| | CCR7 | G043H7 | BV785 | 407 | 780/60 | 1:40 | BioLegend |
| **$CD4^+$ and $CD8^+$ T cell homeostasis** (~1 million PBMCs) | CD3 | UCHT1 | FITC | 488 | 530/30 | 1:25 | BD Pharmingen |
| | CCR10 | 1B5 | PE | 488 | 575/26 | 1:1600 | BD Pharmingen |
| | CD45RA | HI100 | PercP-Cy5.5 | 488 | 695/40 | 1:80 | BioLegend |
| | CD56 | HCD56 | PE-Cy7 | 488 | 780/60 | 1:80 | BioLegend |
| | Fixable Viability dye | na | eFluor660 | 633 | 660/20 | 1:1000 | eBioscience |
| | CD4 | RPA-T4 | APC-Cy7 | 633 | 780/60 | 1:80 | BioLegend |
| | CCR4 | L291H4 | BV421 | 407 | 450/50 | 1:80 | BioLegend |
| | CD45 | HI30 | BV510 | 407 | 525/50 | 1:80 | BioLegend |
| | CCR6 | G034E3 | BV650 | 407 | 660/20 | 1:80 | BioLegend |
| | CxCR3 | G025H7 | BV711 | 407 | 710/50 | 1:80 | BioLegend |
| | CCR7 | G043H7 | BV785 | 407 | 780/60 | 1:40 | BioLegend |
| **Regulatory T cells (Tregs)** (~1 million PBMCs) | CD39 | A1 | FITC | 488 | 530/30 | 1:80 | BioLegend |
| | FoxP3 | PCH101 | PE | 488 | 575/26 | 1:20 | eBioscience |
| | CD45RA | HI100 | PercP-Cy5.5 | 488 | 695/40 | 1:80 | BD Pharmigen |
| | HLA-DR | L243 | PE-Cy7 | 488 | 780/60 | 1:30 | BioLegend |
| | GARP | 7B11 | APC | 633 | 660/20 | 1:80 | eBioscience |
| | CD4 | RPA-T4 | APC-Cy7 | 633 | 780/60 | 1:80 | BioLegend |
| | CD25 | BC96 | BV421 | 407 | 450/50 | 1:20 | BioLegend |
| | Fixable Viability dye | na | BV510 | 407 | 525/50 | 1:1000 | BioLegend |
| | CD45 | HI30 | BV605 | 407 | 610/20 | 1:40 | BioLegend |
| | CD3 | OKT3 | BV650 | 407 | 660/20 | 1:40 | BioLegend |
| | CD127 | A019D5 | BV711 | 407 | 710/50 | 1:20 | BioLegend |
| | CD73 | AD2 | BV785 | 407 | 780/60 | 1:80 | BioLegend |
| **Inhibitory Receptors NK/NKT cells** (~1 million PBMCs) | NKG2A | REA110 | VioBright FITC | 488 | 530/30 | 1:100 | MACS Miltenyi |
| | KIR2DL2/3 | DX27 | PercP-Cy5.5 | 488 | 695/40 | 3:50 | BioLegend |
| | CD57 | HNK-1 | PE-Cy7 | 488 | 780/60 | 1:25 | BioLegend |
| | Fixable Viability dye | na | eFluor660 | 633 | 660/20 | 1:1000 | eBioscience |
| | KIR3DL1 | DX9 | APC-Fire750 | 633 | 780/60 | 1:25 | BioLegend |
| | KLRG1 | 14C2A07 | BV421 | 407 | 450/50 | 3:50 | BioLegend |
| | CD3 | UCHT1 | V500 | 407 | 525/50 | 1:25 | BD Horizon |
| | CD45 | HI30 | BV605 | 407 | 610/20 | 1:25 | BioLegend |
| | CD27 | L128 | BV650 | 407 | 660/20 | 1:25 | BD Horizon |
| | CD56 | NCAM16.2 | BV786 | 407 | 780/60 | 1:25 | BD Horizon |

[1] In a final volume of 100 µL; na, not applicable

**eTable 13 (cont.)**

| Panel | Target | Clone | Fluorochrome | Excitation laser | Detector | Dilution[1] | Company |
|---|---|---|---|---|---|---|---|
| **Activating Receptors NK/NKT cells** (~1 million PBMCs) | NKp44 | P44-8 | PE | 488 | 575/26 | 1:25 | BioLegend |
| | NKp46 | .9E2 | PerCP eFluor711 | 488 | 695/40 | 1:50 | eBioscience |
| | CD57 | HNK-1 | PE-Cy7 | 488 | 780/60 | 1:25 | BioLegend |
| | Fixable Viability dye | na | eFluor660 | 633 | 660/20 | 1:1000 | eBioscience |
| | NKG2D | 1D11 | APC-Cy7 | 633 | 780/60 | 1:25 | BioLegend |
| | CD3 | UCHT1 | V500 | 407 | 525/50 | 1:25 | BD Horizon |
| | CD45 | HI30 | BV605 | 407 | 610/20 | 1:25 | BioLegend |
| | CD27 | L128 | BV650 | 407 | 660/20 | 1:25 | BD Horizon |
| | NKp30 | P30-15 | BV711 | 407 | 710/50 | 1:25 | BioLegend |
| | CD56 | NCAM16.2 | BV786 | 407 | 780/60 | 1:25 | BD Horizon |

[1] In a final volume of 100 µL; na, not applicable

**eTable 14.** Single joint (sj) and DbetaJbeta (DJβ) T cell receptor excision circles (TRECs) and CD3 primers sequences and quantity used in the nested-PCR.

| Primer | | Sequence (5'-3') | Final concentration | Company |
|---|---|---|---|---|
| sjTRECs | OUT 3’ | ACATTTGCTCCGTGGTCTGT | 1 μM | STABVIDA |
| | OUT 5’ | CTCTCCTATCTCTGCTCTGAA | 1 μM | STABVIDA |
| | IN 3’ | GTGCTGGCATCAGAGTGTGT | 1,4 μM | STABVIDA |
| | IN 5’ | TGATGCCACATCCCTTTCAA | 1,4 μM | STABVIDA |
| DJβTREC | OUT 3' | CTCATCTGGGCCTGTCCTTGT | 1 μM | STABVIDA |
| | IN 3’ | TGACCCAGGAGGAAAGAAG | 0,28 μM | STABVIDA |
| | 1.1 - OUT 5' | AACCTAGGACCCTGTGGATG | 1 μM | STABVIDA |
| | 1.1 - IN 5' | TGTCCTCCATCCTAGCCAGG | 0,28 μM | STABVIDA |
| | 1.2 - OUT 5' | CTCTCTATGCCTTCAATGTG | 1 μM | STABVIDA |
| | 1.2 - IN 5' | TCCGTCACAGGGAAAAGTGG | 0,28 μM | STABVIDA |
| | 1.3 - OUT 5' | AAGGGAACACAGAGTACTGGAA | 1 μM | STABVIDA |
| | 1.3 - IN 5' | TCCCAACCTCTGCCTGAAT | 0,28 μM | STABVIDA |
| | 1.4 - OUT 5' | TGGACTTGGGGAGGCAGGA | 1 μM | STABVIDA |
| | 1.4 - IN 5' | AGGAGTGGAAGGCAGCAGGT | 0,28 μM | STABVIDA |
| | 1.5 - OUT 5' | GAAACTGAGAACACAGCCAAGAA | 1 μM | STABVIDA |
| | 1.5 - IN 5' | CTCATAAAATGTGGGTCAGTGGA | 0,28 μM | STABVIDA |
| | 1.6 - OUT 5' | ATCCTCCCTCTTATGTGCATGG | 1 μM | STABVIDA |
| | 1.6 - IN 5' | TGAATCCAGGCAGAGAAAGG | 0,28 μM | STABVIDA |
| CD3γ | OUT 3’ | AGCTCTGAAGTAGGGAACATAT | 1 μM | STABVIDA |
| | OUT 5’ | ACTGACATGGAACAGGGGAA | 1 μM | STABVIDA |
| | IN 3’ | CCTCTCTTCAGCCATTTAAGTA | 1,4 μM | STABVIDA |
| | IN 5’ | GGCTATCATTCTTCTTCAAGGTA | 1,4 μM | STABVIDA |
